# PIAS1 loss and INHBA gain define a TGFβ-driven EMT state in OSCC progression

**DOI:** 10.64898/2026.09.18.752768

**Authors:** Ayan Chanda, Parisa Ghahremanifard, Jinsu An, Rohit Arora, Anusi Sarkar, Angela M. Y. Chan, Doha Itani, Steven C. Nakoneshny, T. Wayne Matthews, Shamir Chandarana, Robert D. Hart, Joseph C. Dort, Martin D. Hyrcza, Shirin Bonni, Pinaki Bose

**Affiliations:** Department of Biochemistry and Molecular Biology, Cumming School of Medicine, University of Calgary, Calgary, AB, Canada; Charbonneau Cancer Institute, Arthur J.E. Child Comprehensive Cancer Centre, Cumming School of Medicine, University of Calgary, Calgary, AB, Canada; Department for Biomedical Informatics, Harvard Medical School, Boston, MA, USA; Department of Pathology and Laboratory Medicine, Cumming School of Medicine, University of Calgary, Calgary, AB, Canada; Cancer Translational Research Core, Arthur J.E. Child Comprehensive Cancer Centre, Alberta Health Services, Calgary, AB, Canada; Molecular Diagnostics and Cytogenetics, Saint John Regional Hospital, Horizon Health Network, Saint John, NB, Canada; Ohlson Research Initiative, Arnie Charbonneau Cancer Institute, Cumming School of Medicine, University of Calgary, Calgary, AB, Canada; Department of Surgery, Section of Otolaryngology – Head & Neck Surgery, Cumming School of Medicine, University of Calgary, Calgary, AB, Canada; Department of Oncology, Cumming School of Medicine, University of Calgary, Calgary, AB, Canada

**Author notes:** These authors contributed equally to this work (Ayan Chanda, Parisa Ghahremanifard). Authors are listed alphabetically. Corresponding author: Pinaki Bose –. **Competing Interests:** The authors declare no competing interests.

## Abstract

Oral squamous cell carcinoma (OSCC), a highly aggressive subtype of head and neck cancer, remains a major clinical challenge with limited therapeutic options and poor survival outcomes. The Protein Inhibitor of Activated STAT1 (PIAS1), a SUMO E3 ligase involved in transcriptional regulation, DNA repair, and epithelial–mesenchymal transition (EMT), exhibits cancer type–specific roles, but its function in OSCC has not been well characterized. Here, we investigated the molecular and functional role of PIAS1 in OSCC progression and prognosis. All patient cohorts analysed here comprise HPV-negative disease. Analysis across multiple cohorts revealed that PIAS1 mRNA expression is significantly lower in tumor tissues than in normal oral cavity squamous epithelium, including in a cohort of 60 patient-matched tumor and normal pairs. Within the tumor compartment, protein abundance of PIAS1 correlated positively with patient survival. Functional assays using OSCC cell lines demonstrated that PIAS1 acts in a SUMO E3-dependent manner to suppress TGFβ-induced migration and invasion. Transcriptomic profiling of PIAS1-depleted cells revealed alterations in expression of genes with relevance in TGFβ signaling and pro-invasive cellular responses. Further analyses showed that 46 of these differentially altered hits intersected with The Cancer Genome Atlas (TCGA)-OSCC datasets, where four of these genes, including INHBA, associated with cell stemness-pathway. Spatial analysis of the mRNA abundance of INHBA within OSCC-derived surgical specimens showed increased abundance at the invasive front, and positively associated with aggressive tumor phenotypes, poor patients’ outcomes, and resistance to immunotherapy. Single-cell RNA-seq analyses of INHBA revealed increased levels within tumor cells and cancer-associated fibroblasts (CAFs), suggesting potential tumor-intrinsic and extrinsic (tumor microenvironment) sources and hence actions. Collectively, our findings establish PIAS1 as a tumor suppressor in OSCC that restrains TGFβ-driven aggressive behaviour. Furthermore, we identify INHBA as a potential therapeutic target, among other genes characterized by spatially restricted, invasive tumor phenotypes.

**Graphical Abstract:** 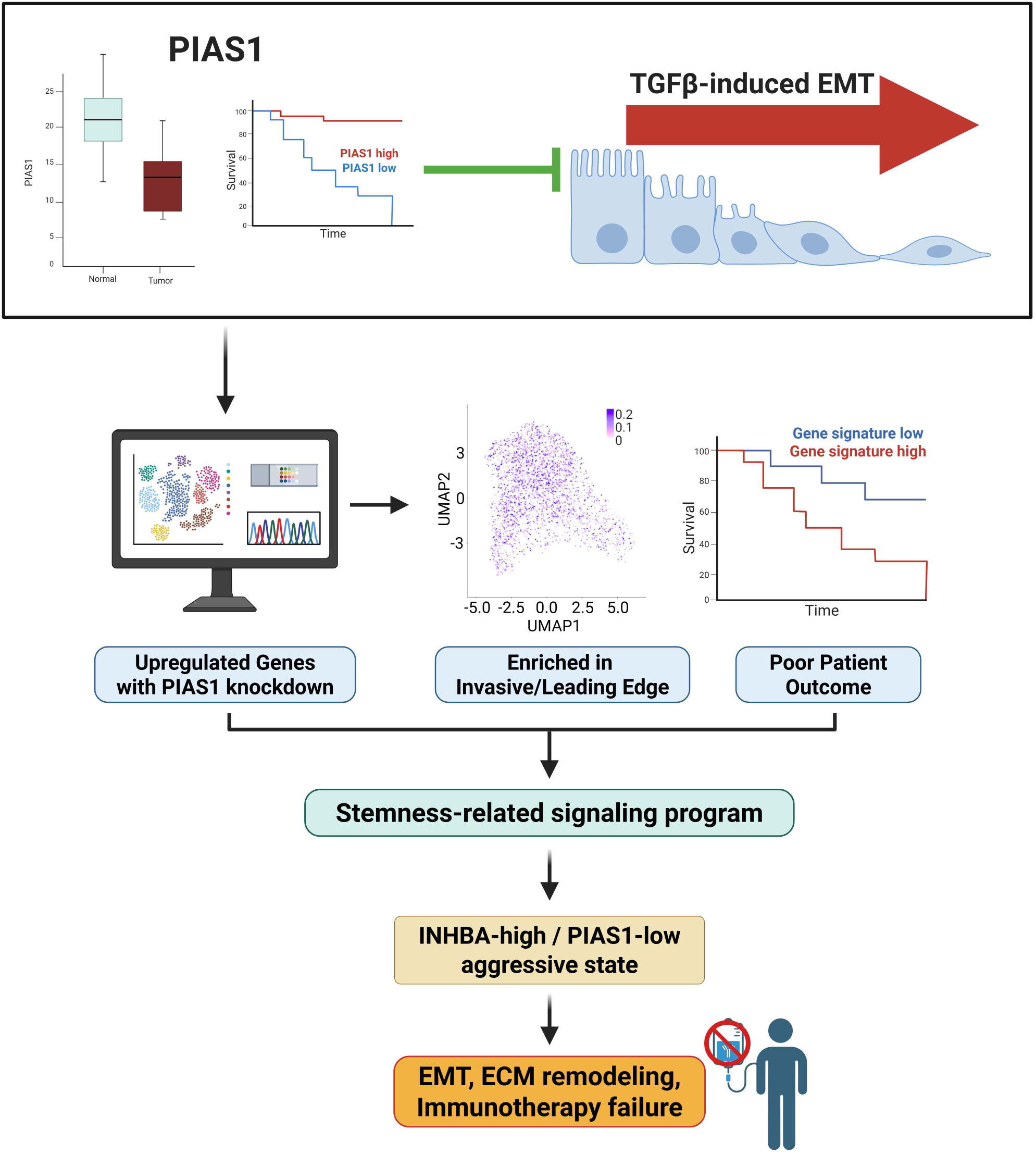

## Introduction

OSCC is the most common subtype of non-cutaneous head and neck cancer that is associated with high morbidity and mortality (1). Despite advances in surgical and non-surgical therapeutic approaches, 5-year survival for OSCC patients remains at ∼50% (2, 3). Approximately 40% of OSCC patients present with cervical lymph-node metastasis (LNM) at diagnosis, reflecting the aggressive nature of cancer cells that escape from the primary tumor site. The presence of LNM significantly worsens prognosis and contributes to poor clinical outcomes (4). Although LNM is the main reason for mortality among OSCC cancer patients, the cellular mechanisms driving metastatic outgrowth in OSCC patients remains largely unknown (5). Identifying and characterizing novel factors that regulate oral cancer progression and LNM are critical in developing targeted therapies and improving survival rates.

Epithelial-mesenchymal transition (EMT) is an important cellular process that contributes to tumor progression and metastasis (6, 7). During EMT, epithelial cells lose their apical-basal polarity and adhesion properties, transitioning into a mesenchymal-like state with increased motility and invasive capacity (8). Cells undergoing EMT secrete proteases to degrade the extracellular matrix (ECM), therefore allowing easy movement and invasion, facilitating spread loco-regionally and to distant sites (7, 9). A key signaling axis that induces EMT is the transforming growth factor-beta (TGFβ)-signaling pathway, which drives increased or decreased expression of sets of genes that control EMT (10, 11). TGFβ-induced EMT has been suggested to promote OSCC progression (12). Consequently, identifying factors and cellular processes that can inhibit TGFβ-induced EMT represents a critical avenue of investigation in OSCC research.

Sumoylation is a post-translational modification involving the covalent attachment of small ubiquitin-like modifiers (SUMOs) to lysine residues of target proteins through a multi-step enzymatic process (13). Among components of the SUMO cascade, the SUMO E3 ligases are critical regulators of substrate recognition and modification (14). Protein inhibitor of activated STAT1 (PIAS1) is a SUMO E3 ligase that has many substrates, including several tumor promoters and suppressors (14). PIAS1 can regulate key cellular functions such as development, differentiation, apoptosis, and senescence (13, 14). Notably, PIAS1 has been implicated in suppression of transcription of TGFβ-responsive genes and inhibition of EMT in mammary epithelial and breast cancer cells (15–18). These reports raise two key questions in OSCC: 1) whether PIAS1 acts in a SUMO E3 ligase-dependent manner to suppress TGFβ-responsive genes involved in cell plasticity, including EMT, and 2) whether PIAS1 expression or activity is mediated by downstream oncogenic mechanisms. Investigating whether downregulation of PIAS1 contributes to tumor progression by enhancing key oncogenic pathways, including those driving invasion and metastasis, could uncover novel therapeutic targets and enhance its potential as a prognostic biomarker in OSCC.

In this study, we find that compared to normal oral cavity squamous epithelium (OCSE), there is a reduction in mRNA and protein abundance of PIAS1 in OSCC tumor samples. Importantly, our tissue-microarray analyses indicate that higher protein abundance of PIAS1 in OSCC correlates with improved survival outcomes. Functionally, we find that PIAS1 acts in a SUMO E3 ligase-dependent manner to suppress TGFβ-induced migratory behavior and invasive growth of OSCC cell lines in two- and three-dimensional cultures, respectively. Consistently, spatial transcriptomic profiling and RNA sequencing data revealed a PIAS1-regulated gene signature, including the targetable oncogenic effector INHBA, which we uncovered to positively associate with aggressive tumor phenotypes in OSCC. Collectively, these findings suggest that PIAS1 may play a critical tumor suppressive role in OSCC with prognostic value in OSCC, and that its newly revealed downstream oncogenic effectors may serve as potential therapeutic targets in OSCC and other cancers.

## Material and methods

### In silico Data Analysis

Publicly available datasets from The Cancer Genome Atlas (TCGA) were utilized to investigate PIAS1 expression and its downstream effects in HPV-negative OSCC samples. Raw/normalized expression values and clinical information were downloaded from UCSC Xena (19). Samples were filtered to include cases that are within the oral cavity that are HPV-negative. DESeq2-based differential gene expression analyses comparison of lowest (n=92) versus highest (n=92) tertiles of PIAS1 expression. Differentially expressed genes (DEGs) were defined by a log2-fold change threshold (± 0.58) with an adjusted p-value <0.05. The identified DEGs were visualized using principal component analysis (PCA) and volcano plots. Ingenuity Pathway Analysis (IPA; Qiagen, CA, USA) was employed to identify DEGs-impacted canonical pathways and upstream regulators. In parallel, single-sample gene set enrichment analysis (ssGSEA) was conducted to assess EMT-related gene signatures. For the PIAS1-associated signatures derived from the CAL33 and TCGA overlap, ssGSEA scores were computed with each gene weighted by its log2 fold change in the CAL33 PIAS1i versus control comparison, so that genes with larger differential expression contributed proportionally more to the sample-level score. The TGFβ and EMT gene sets were obtained from the MSigDB C2 collection, specifically removing the common genes associated with the c-ECM-UP signature (20). In addition to TCGA, the mRNA and protein abundance of PIAS1 were analyzed using the Clinical Proteomic Tumor Analysis Consortium (CPTAC) head and neck squamous cell carcinoma (HNSCC) patient cohort data and were assessed for association with patient survival (21). Matched tumor (n=108) and normal samples (n=63) were analyzed to evaluate the mRNA and protein abundance of PIAS1 in tumor and normal-derived samples, and the correlation between mRNA and protein abundance in tumor and normal samples was also assessed. Datasets from a cohort (Calgary) of in-house matched tumor and normal samples (n=60) were used to validate the findings obtained from TCGA and CPTAC sources (Suppl. Table 1). Statistical comparisons were performed using non-parametric Wilcoxon rank-sum tests, with p-values <0.05 considered significant. Gene expression data for normal human tissue samples from the Genotype-Tissue Expression (GTEx) project (22) were downloaded from the GTEx portal.

A single-cell RNAseq dataset was obtained from Puram *et al.* (GSE103322, (23)) and contained ∼6000 cells from 18 OSCC patients. The Seurat package (version 5.0.1) was used for analysis. After excluding samples from metastatic sites, we retained a total of 4486 cells (1706 malignant and 2780 non-malignant). We carried out quality control steps that included filtering out cells with low gene counts and normalizing the retained data to minimize effects of technical variations. Following clustering and cell-type annotation, the relative mRNA abundance of specific genes were evaluated across different cell populations within the tumor microenvironment (TME).

Single-cell RNA-seq datasets were assembled from four published OSCC cohorts and restricted to primary OSCC tumors only (24–27). Global quality control filtering was applied using nCount_RNA ≥ 400, nFeature_RNA ≥ 250, and percent.mt ≤ 15, followed by per-sample doublet removal with scDblFinder (28). Only cells passing quality control and classified as singlets were retained for downstream reference construction. Each dataset was then processed independently using a standard Seurat workflow, including SCTransform normalization, principal component analysis, shared nearest-neighbor graph construction, and Louvain clustering.

Gene expression profiles of responders and non-responders to immune checkpoint blockade were obtained from seven independent cohorts, comprising a total of 335 patients across melanoma, urothelial carcinoma, and pancreatic adenocarcinoma (29–35). Treatment regimens differed across cohorts and included anti-PD-1 monotherapy, anti-PD-L1 monotherapy, combined anti-PD-1 and anti-CTLA-4, and an anti-CD40 agonist with or without anti-PD-1. No head and neck cohort with matched transcriptomic and response data was available. For immunotherapy response analyses, INHBA-PIAS1 scores were calculated within each cohort after dataset-specific normalization and standardization to avoid cross-cohort scale effects. Patients were classified as responders or non-responders according to the clinical response labels provided by each study. Score differences between response groups were assessed using Wilcoxon rank-sum tests, and cohort-combined analyses were performed after within-cohort scaling.

Within each dataset, INHBA and PIAS1 values were standardized to z-scores to place both markers on a comparable scale. The INHBA-PIAS1 score was calculated for each sample or spatial spot as z(INHBA) minus z(PIAS1), such that higher values represented relatively higher INHBA abundance and relatively lower PIAS1 abundance. For group-wise comparisons, samples were stratified into high-score and low-score groups using tertile cutoff unless otherwise specified. For spatial analyses, the same score was calculated at the spot level using normalized spatial expression values.

### Gene set enrichment and pathway scoring

Gene set enrichment analyses were performed to determine whether tumors with high INHBA-PIAS1 scores were enriched for extracellular matrix remodeling, TGFβ-associated, and EMT-related programs. Samples were ranked according to differential expression or differential abundance between high-score and low-score groups. Enrichment was tested using curated gene sets including, Foroutan-TGFβ regulated EMT Down Signature, TGFβ-upregulated gene signatures, and EMT-associated gene sets from MSigDB and the cancer-associated extracellular matrix upregulated signature (c-ECM-UP) signature from Chakravarthy et al. Normalized enrichment scores and multiple-testing-adjusted q-values were reported. For single-sample or spot-level pathway analyses, module scores were calculated using normalized expression matrices, and associations with the INHBA-PIAS1 score were evaluated using Spearman correlation or Wilcoxon rank-sum tests, as appropriate.

### OSCC Tissue MicroArray (TMA)-Cohort

The Ohlson Research Initiative (ORI; University of Calgary) OSCC cohort comprises histologically confirmed, surgically-resected treatment naive tumors from 175 patients diagnosed between 2009 and 2013 (Suppl. Table 2) (36). The median age and median duration of follow-up were 62.5 years and 5.8 years, respectively. The collection of patient materials and clinical data followed Tri-council Policy Statement for Research with Human Subjects’ guidelines (Canada). The study was conducted in accordance with the reporting recommendations for tumor marker prognostic studies (REMARK) guidelines (37), and was approved by the Health Research Ethics Board of Alberta (HREBA).

### Fluorescence immunohistochemistry (IHC)

Fluorescence IHC was performed on TMAs. To account for variability in staining across multiple TMA slides, each TMA was stained alongside an on-slide reference control containing cell lines with increasing protein abundance of PIAS1 (17) that was used to normalize the relative PIAS1-signal intensities across the entire study. The on-slide protein abundance of PIAS1 in control cells were analyzed using the same digital image analysis pipeline as the patient TMA cores. A reference TMA was selected to generate a standard curve for normalization and to ensure consistency in PIAS1 protein quantification across all TMAs (17). Antigen retrieval was performed using citrate-based target retrieval solution (Dako, catalog number S1699), with heating to 121°C for three minutes in a decloaking chamber (Biocare Medical). A rabbit monoclonal anti-PIAS1 antibody (clone EPR2580(2), 1:2000, Abcam) and a mouse monoclonal anti-pan-cytokeratin antibody (clone AE1/AE3, 1:100, Dako) were used as primary antibodies. In addition, isotype control antibodies at equivalent concentrations to the respective primary antibodies were included as negative controls. Secondary antibodies included anti-rabbit and anti-mouse EnVision+ (K4011, Dako), and signal amplification was achieved using TSA-Plus Cy5 reagents (PerkinElmer). Nuclei were counterstained with DAPI (Thermo Fisher Scientific, catalog number D1306). After immunostaining, slides were coverslipped using ProLong Gold anti-fade mounting medium (Thermo Fisher Scientific, catalog number P36934) and stored at 4°C until imaging. In-house immunoblotting and control tissue immunostaining were previously employed to confirm the target specificity of the primary antibody used in this study (17).

### Quantitative image analysis

Digital image analysis involved using an Aperio ScanScope Fl. Identical image acquisition parameters were applied to all labelled slides. To determine the mean staining intensity, an algorithm was designed within the HALO image analysis software platform (version 2.0.1145.14, Indica Labs) (17). A tumor-specific mask was generated to distinguish the cancer cells from surrounding stromal tissue by thresholding the pan-cytokeratin images, a stromal-specific mask was generated as the inverse of the tumor-specific mask. Thresholding levels were verified and adjusted, if necessary, by spot-checking a small sample of images to determine an optimal threshold value. All images were then processed using this optimal threshold value and all subsequent image manipulations involved only image information in a masked area. Unusable areas such as folded or necrotic tissue were manually cropped. Criteria for inclusion of TMA cores in the analyses included: 1) 50% of imaged-tissue was well focused with no overlap, and 2) the selected TMA core comprised of more than 200 cells. After review and image analysis validation, data from patients that had usable results were used for subsequent statistical analysis.

### Cell culture and transfection

CMV-based plasmids containing cDNA encoding FLAG-tagged un-mutated wild type PIAS1 (WT), or PIAS1 in which Cysteine 350 was converted to serine to produce a SUMO E3 ligase inactive (CS) version of PIAS1, as well as a U6-CMV-based plasmid containing either no hairpin or a short hairpin RNA (shRNA) against PIAS1 (PIAS1i) downstream of the U6 promoter, together with a cDNA encoding enhanced green fluorescent protein (GFP) downstream of the CMV promoter, as described previously (16). Expansion of the plasmids were performed by transforming Escherichia coli DH5α^TM^ (Invitrogen, Canada) using the heat-shock method. The plasmids were extracted from positive colonies using Qiaprep Kit (Qiagen, USA).

The human oral tongue-derived SCC CAL33 cell line was cultured in Dulbecco’s modified Eagle’s medium with high glucose and L-glutamine (DMEM) (Invitrogen, Canada)-containing 10% fetal bovine serum (FBS, Invitrogen, Canada), 1% penicillin-streptomycin (Thermo Fisher Scientific) and 1% MEM Non-Essential Amino Acids (Thermo Fisher Scientific). CAL33 cell cultures, which were kept at 37°C in a 5% CO_2_-containing incubator, were passaged every 3-4 days. CAL33 cell cultures were routinely tested for mycoplasma using the MycoAlert® Mycoplasma Detection Kit (Lonza, NY, USA) testing to ensure negativity. The Lipofectamine 3000 or RNAiMAX reagent (Invitrogen, Canada) was used to introduce genetic material into these SCC cells. Recombinant human TGF-β1 (R&D Systems, catalog number 240-B) was reconstituted according to the manufacturer’s instructions and used at a final concentration of 100 pM (approximately 2.5 ng/mL) in all experiments.

### Scratch Assay

Transfected CAL33 cells were seeded in a 12-well plate for 24h to reach near full-confluency in complete medium, followed by 24h serum starvation by culturing in 0.2% FBS-containing medium. Scratch assay was performed by gliding a 200 μL pipette tip across the cell surface to produce a vertical line connecting the 12 and the 6 positions of a 12-hour clock. After PBS-based removal of floating cells and wash of adherent cells, cells in the 12-well tissue culture plates were incubated in 0.2% FBS-medium with or without TGFβ, and were incubated for 30h at 37°C in the 5% CO2 humidified cell incubator (38, 39). In each well, five images of the scratch were captured at a 4X objective (Olympus CKX53SF) at times 0h and 30h post-incubation. ImageJ (National Institutes of Health, USA) was used to obtain three measurements per image along different locations within the cell-free scratch region, for a total of 15 values per well/experimental condition, the mean of which was used for downstream analysis. Cell migration is defined as relative wound closure distance between 0 and 30 h per condition. Three biological replicates were carried out per each experiment (39).

### Transwell Invasion Assay

Transwell invasion assay was performed using polycarbonate filters (24-well inserts, pore size 8 μm; BD Biosciences, Canada). First, each well and insert membrane were equilibrated by adding 500 μL of serum-free DMEM and incubating for 2h at 37°C incubator. After removing the equilibration media, the top surface of each insert was coated with 50 ul of 3% Matrigel (Corning Incorporated, USA) and incubated at 37°C incubator for 1h to allow the Matrigel to solidify. 2 × 10^5^ overnight serum-starved CAL33 cells, resuspended in 500 μL of serum-free medium were seeded on the upper side of the Matrigel-coated filter. 500 μL of complete medium without or with TGFβ was added to the lower chamber. After 24h of incubation at 37°C in 5% CO2 incubator, non-adherent cells on the upper chamber were removed by washing with PBS three times. In the second wash using a cotton tip applicator, adherent cells on the upper surface of the membrane were gently scraped off. Cells that had invaded onto the underside of the inserts were fixed with 100% methanol for 10min at -20 °C, and stained with 0.5% crystal violet dye (EMD Millipore, Canada) at room temperature for 1h. Images were captured from 5 random locations at 10× objective (Olympus CKX53SF) for each experimental condition. Invaded cells were counted and summed for each condition. Each experiment was repeated three independent times and average biological replicates were subjected to statistical analysis (16, 39).

### Three-dimensional culture

Three-dimensional (3D) cultures of cells were prepared in 96-well flat-bottom, ultra-low attachment plates (BD Biosciences, ON, Canada). Each well of the 96-well dish was precoated with 50 μL of ice-cooled 3 mg/ml final concentration growth-factor reduced Matrigel (Corning Incorporated, USA) mixed with antibiotic–penicillin/streptomycin (Thermo Fisher Scientific)-containing full growth medium. The Matrigel bed was allowed to solidify by incubating for 1h at 37 °C in 5% CO_2_ containing tissue culture incubator. Approximately 300 cells per well of 96-well dish were resuspended in 50 μl of 5 mg/ml ice-cooled Matrigel in complete growth medium and layered on top of the Matrigel bed and allowed to solidify by incubating at 37 °C for 1h in 5% CO_2_ containing tissue culture incubator. 50ul of complete medium was layered on top after 1 h. The next day and every third day, the culture received complete media alone or together with 100pM TGFβ. Isolated cells divided and formed multicellular structures referred to as organoids. Images of six representative 8-day old organoids from each well were captured at 40× objective (Olympus CKX53SF) followed by an assessment of growth phenotypes (spherical or non-deformed vs. deformed organoids). Three independent replicates of each experiment were performed and average percentage of spherical organoids were calculated.

### RNAseq analysis

Triplicate adherent cultures of CAL33 cells were transfected with a control or PIAS1i plasmid. 2 days following start of the transfections, each of the cell populations were subjected to RNeasy Plus Minikit RNA extraction (Qiagen, ON, Canada). The quality/integrity of the extracted RNA was assessed using the Qubit RNA IQ Assay Kit (Thermo Fisher Scientific) with a Qubit 4 Fluorometer (Invitrogen, Canada) and expressed as RNA Integrity Number (RIN) scores. RNA samples with RIN scores of 7 and higher were used to generate RNAseq libraries using the NEBNext Ultra II Directional RNA Library Prep Kit for Illumina (NEB, ON, Canada), and then sequenced on the NextSeq platform (Illumina, CA, USA) to generate single-end 50 bp reads at a target depth of 20-30 million reads per sample. Raw read quality was assessed with FastQC and summarised with MultiQC. Adapters and low-quality bases were removed with Trim Galore using default parameters. Surviving reads were aligned to the human reference genome GRCh38 with STAR with GENCODE release 47 annotation. Differential expression analysis was performed on the filtered raw counts using the DESeq2 Bioconductor package (40) to compare gene expression profiles in PIAS1i and control cells. Genes with a log2 fold-change threshold of ±0.58 and an adjusted p-value <0.05 were considered as differentially expressed genes (DEGs). DEGs were next analyzed using QIAGEN IPA to compare alterations in canonical signaling pathways and their predicted upstream regulators between the PIAS1 knockdown and control conditions. We then compared this study’s identified DEGs to those retrieved from the TCGA portal. Overlapping genes were displayed using a Venn diagram, followed by IPA-based pathway analysis and assessing the literature for possible relevance to OSCC tumorigenesis and progression. An extensive literature review was performed to evaluate the roles (if any) of the DEGs in regulating OSCC tumorigenesis and progression.

### Spatial Transcriptomics Analysis

Spatial transcriptomics data were generated using the Visium platform (10x Genomics) on 12 surgically resected, fresh-frozen OSCC tumors obtained from 10 patients. To provide histopathological context for spatially resolved molecular analyses, the study pathologist (MH) manually reviewed the corresponding H&E-stained sections and annotated major tumor compartments, including the tumor core (TC), transitory zone, and leading edge (LE). Spatial transcriptomic data were processed and analyzed using the Seurat package. To resolve the cellular composition of individual spatial capture location, Visium data were deconvolved using CARD (41) with an independently annotated OSCC single-cell RNA-sequencing reference dataset (GSE103322). CARD-derived cell-type abundance scores were incorporated into the spatial Seurat object and used to characterize the distribution of malignant, stromal, and immune populations across the tumor microenvironment. These estimates were subsequently used to define cancer- and fibroblast-enriched spatial compartments and to investigate relationships between the PIAS1–INHBA transcriptional state and tumor microenvironment composition.

Gene-expression programs identified from bulk transcriptomic analyses were projected onto the spatial transcriptomic data using the Seurat AddModuleScore() function (32). Additional curated transcriptional programs representing TGF-β signaling, epithelial–mesenchymal transition, CAF/ECM remodeling, myofibroblast contractility, hypoxia, cytotoxic T-cell activity, tumor-infiltrating lymphocytes, and antigen-presentation programs were similarly evaluated at the spatial-location level.

To evaluate the spatial distribution of the PIAS1–INHBA state, spatial locations were classified according to PIAS1 and INHBA expression-derived states, including a PIAS1-low/INHBA-high state of interest and a reciprocal PIAS1-high/INHBA-low state. The proportion of PIAS1-low/INHBA-high spatial locations was quantified within pathologist-defined tumor compartments on a per-sample basis. Comparisons between spatial regions were performed using paired non-parametric tests, with P-values adjusted for multiple comparisons using the Benjamini–Hochberg method where applicable.

Associations between the continuous PIAS1–INHBA transcriptional axis and tumor microenvironment composition were assessed independently within each specimen using Spearman rank correlations between the PIAS1–INHBA axis and CARD-derived cell-type abundance scores, including CAF, myofibroblast, T-cell, and myeloid populations. Associations with spatial gene-expression programs were assessed similarly. Sample-specific correlation coefficients were summarized across the cohort, and the consistency of associations across samples was evaluated using Wilcoxon signed-rank tests with Benjamini–Hochberg correction for multiple testing.

To compare gene expression levels across different spatial regions, we performed pairwise non-parametric Wilcoxon rank-sum tests, evaluating differentially expressed genes (DEGs) between tumor core (TC), leading edge (LE), and transitory zone regions. P-values were adjusted to correct for multiple testing using the Benjamini-Hochberg method, with a significance threshold of adjusted p-value (false discovery rate; FDR) < 0.05.

To assess transcriptional dynamics and cellular state transitions, we applied RNA velocity analysis using the velocyto command-line tool, following established workflows (42). Briefly, spliced and unspliced mRNA counts were extracted from Visium spatial transcriptomic BAM files. Cell-type-specific velocity vectors were computed to infer potential cell-state transitions within the TME. The velocity vectors were projected onto the spatial transcriptomics UMAP space, providing insight into transcriptional trajectories and potential lineage differentiation pathways within OSCC tumor compartments. To model the effects of specific gene perturbations on spatial cell fate decisions, we utilized Dynamo (Version 1.2.0, Python package), an advanced RNA velocity-based dynamical modeling tool. Specifically, perturbation simulation of INHBA was conducted using the dyn.pd.perturbation() function (42) applying a Jv scaling factor of -1000, representing a hypothetical strong downregulation scenario. The resulting perturbation data were integrated into the velocity vector space, allowing for an estimation of spot fate transition probabilities among tumor core, transitory zone, and leading-edge states. To quantitatively assess the impact of INHBA modulation on spatial transitions, we constructed a state transition probability graph using the dyn.pd.state_graph() function in velocity field (vf) mode. This approach enabled a spatially resolved prediction of INHBA-driven cell state alterations, providing a mechanistic link between transcriptional reprogramming and spatial tumor evolution.

### Cell-cell communication analysis

Cancer–CAF ligand–receptor interactions were evaluated from the deconvolved spatial transcriptomic data described above. Curated ligand–receptor pairs and their pathway assignments were taken from the human CellChatDB as distributed with CellChat v2.1.2 (43)); interaction strength was computed directly from the spatial expression matrices as described below rather than by the CellChat permutation-based inference procedure. Cancer-like and CAF-like spatial locations were defined from CARD-derived scores standardized within each sample; spots were classified as cancer-like when the cancer z-score was ≥0 and exceeded the CAF z-score, and as CAF-like when the CAF z-score was ≥0 and exceeded the cancer z-score. Spots were further stratified into PIAS1-low/INHBA-high and PIAS1-high/INHBA-low states, and only samples containing ≥20 spots in each of the four resulting groups were retained. For each sample, normalized expression was averaged within each group, and ligand– receptor support was calculated as mean ligand expression in the sender multiplied by mean receptor expression in the receiver. Cancer-to-CAF and CAF-to-cancer interactions were analyzed separately, and differential interaction strength (ΔLR) was defined as the PIAS1-low/INHBA-high score minus the corresponding PIAS1-high/INHBA-low score.

### Statistical Analysis

Kaplan–Meier survival curves were generated to evaluate differences in patient outcomes across gene expression groups, and survival distributions were compared using the log-rank test. Cox proportional hazards regression analysis was performed to estimate hazard ratios (HR) and corresponding 95% confidence intervals for overall survival (OS), disease-specific survival (DSS), and progression-free interval (PFI) associated with gene set expression.

For comparisons between two groups of bulk RNA-sequencing or spatial transcriptomics samples, the Wilcoxon rank-sum test was used. Biochemical and organoid-based experimental data were analyzed using an unpaired Student’s t-test for two-group comparisons. Experiments with a factorial design, comprising construct (vector, PIAS1i, PIAS1 WT, or PIAS1 CS) and treatment (untreated or TGFβ), were analyzed by two-way analysis of variance (ANOVA) with an interaction term, followed by the Tukey–Kramer post hoc test for pairwise comparisons among groups. The construct-by-treatment interaction term was used to test whether PIAS1 status modifies the response to TGFβ. Statistical analyses were performed using GraphPad Prism version 8.0 (GraphPad Software, San Diego, CA, USA).

Data are presented as mean ± standard error of the mean (SEM). All experiments were independently repeated at least three times. A P value ≤ 0.05 was considered statistically significant. Significance levels are indicated as p < 0.05 (*), p < 0.01 (**), p < 0.001 (***), and p < 0.0001 (****); ns denotes not significant.

## Results

### mRNA and protein abundance of PIAS1 negatively correlate with OSCC progression and prognosis

To investigate if PIAS1 is dysregulated in OSCC, we first asked whether PIAS1 is differentially expressed in tumor vs normal samples using the publicly accessible TCGA transcriptomics datasets for this cancer. Our analyses demonstrated a downregulation of the relative mRNA abundance of PIAS1 in OSCC tumor samples (n=275) compared with OCSE (n=25) (Fig. 1A). Similar results were observed in our in-house cohort of 60 matched OSCC and OCSE samples profiled using RNAseq (Fig. 1B), as well as a cohort of HNSCC curated by the Clinical Proteomic Tumor Analysis Consortium (CPTAC) containing 108 tumor and 63 normal samples (Fig. 1C). Importantly, the PIAS1 mRNA expression and protein abundance data from CPTAC patients revealed a positive correlation (Suppl. Fig. 1A). In addition, we found that the protein abundance of PIAS1 is reduced in tumors compared with normal samples (Fig. 1D). Further analyses of the CPTAC cohort revealed that high PIAS1 protein abundance in tumors was associated with improved overall survival (Figure 1E).

**Figure 1:**
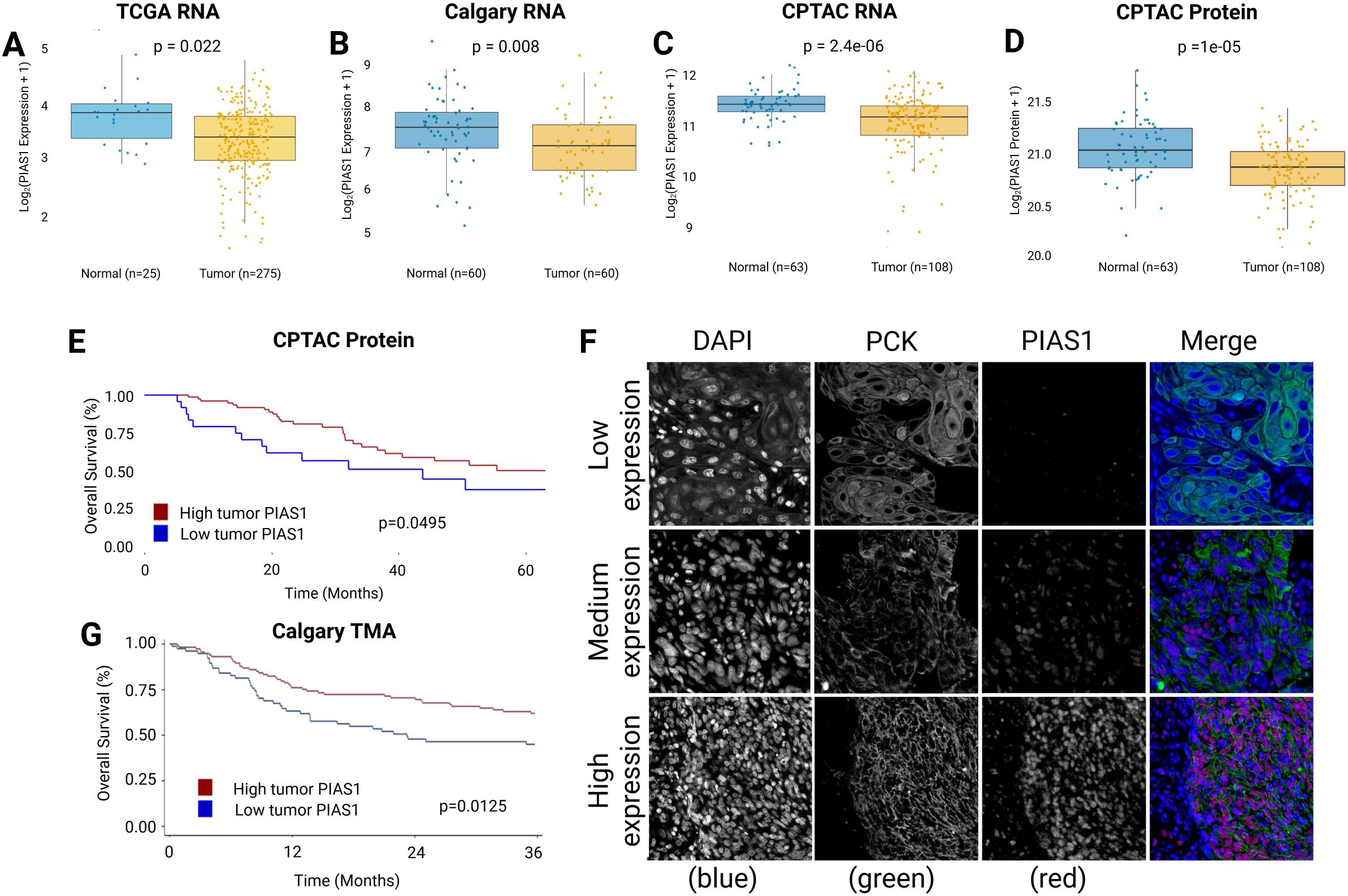
mRNA and protein abundance of PIAS1 are reduced in OSCC tumors and higher PIAS1 is associated with improved patient survival. (A) Box plot showing PIAS1 mRNA expression in OSCC tumor samples (n=275) compared to normal oral cavity squamous epithelium (OCSE; n=25) in the TCGA cohort. (B) Box plot showing PIAS1 mRNA expression in matched OSCC tumor samples (n=60) compared to normal tissues (n=60) in the Calgary cohort. (C) Box plot showing PIAS1 mRNA expression in tumor samples (n=108) compared to normal tissues (n=63) in the CPTAC cohort. (D) Box plot showing PIAS1 protein abundance in tumor samples (n=108) compared to normal tissues (n=63) in the CPTAC cohort. (E) Kaplan-Meier curve for overall survival (OS) based on PIAS1 protein abundance in patients from the CPTAC cohort, dichotomized into high and low tumor PIAS1 groups. (F) Representative immunofluorescence images of OSCC patient samples from the Calgary TMA cohort showing low, medium, and high PIAS1 expression. DAPI-stained nuclei are shown in blue, pan-cytokeratin-positive tumor cells are shown in green, and PIAS1 staining is shown in red. (G) Kaplan-Meier curve for OS based on tumor PIAS1 protein abundance in the Calgary TMA cohort, dichotomized into high and low tumor PIAS1 groups.

To orthogonally validate the prognostic value of PIAS1 in OSCC tumors, we performed quantitative fluorescence immunohistochemistry on TMAs containing primary tumor and matched normal OCSE from the ORI cohort. Here, a PIAS1 antibody was used that has been confirmed to have high specificity and sensitivity in detecting varying protein abundance of PIAS1 in human tissues (17). OSCC patient samples were divided into high and low groups based on the median HALO score that corresponded to the fluorescence intensity of PIAS1 staining in tumor tissue (Fig. 1F). Patients with high PIAS1 tumors were associated with improved overall survival compared with patients with low PIAS1 tumors (Fig. 1G). These findings suggest that increased protein abundance of PIAS1 in tumors contributes to improved prognosis.

### PIAS1 regulates the migration and invasive growth of the OSCC cells

Given the association of PIAS1 with improved patient outcome, we tested its functional relevance in OSCC cells. Surgical margin status refers to the presence or absence of cancer cells at the edge of the excised tissue. A positive margin indicates that cancer cells are present at the resection boundary, suggesting incomplete tumor removal, while conversely, a negative margin implies no cancer cells are detected, indicating likely complete tumor resection (44). Margin positivity is an established marker of worse prognosis in OSCC that is associated with increased rates of locoregional recurrence, reduced disease-free and overall survival rates (44, 45). Residual tumor cells at positive margins, often undergoing EMT, exacerbate the likelihood of disease recurrence and contribute to the aggressive nature of the disease (46, 47). We observed that PIAS1 expression is significantly lower in tumors associated with positive margins as compared to those with negative margins in the TCGA OSCC cohort (Fig. 2A). Further analyses revealed that relative mRNA abundance of PIAS1 correlated positively with a TGFβ-induced “EMT-downregulated” gene signature (20) (Fig. 2B). Together, these analyses support a role for PIAS1 as a potential suppressor of EMT-like features in OSCC.

**Figure 2:**
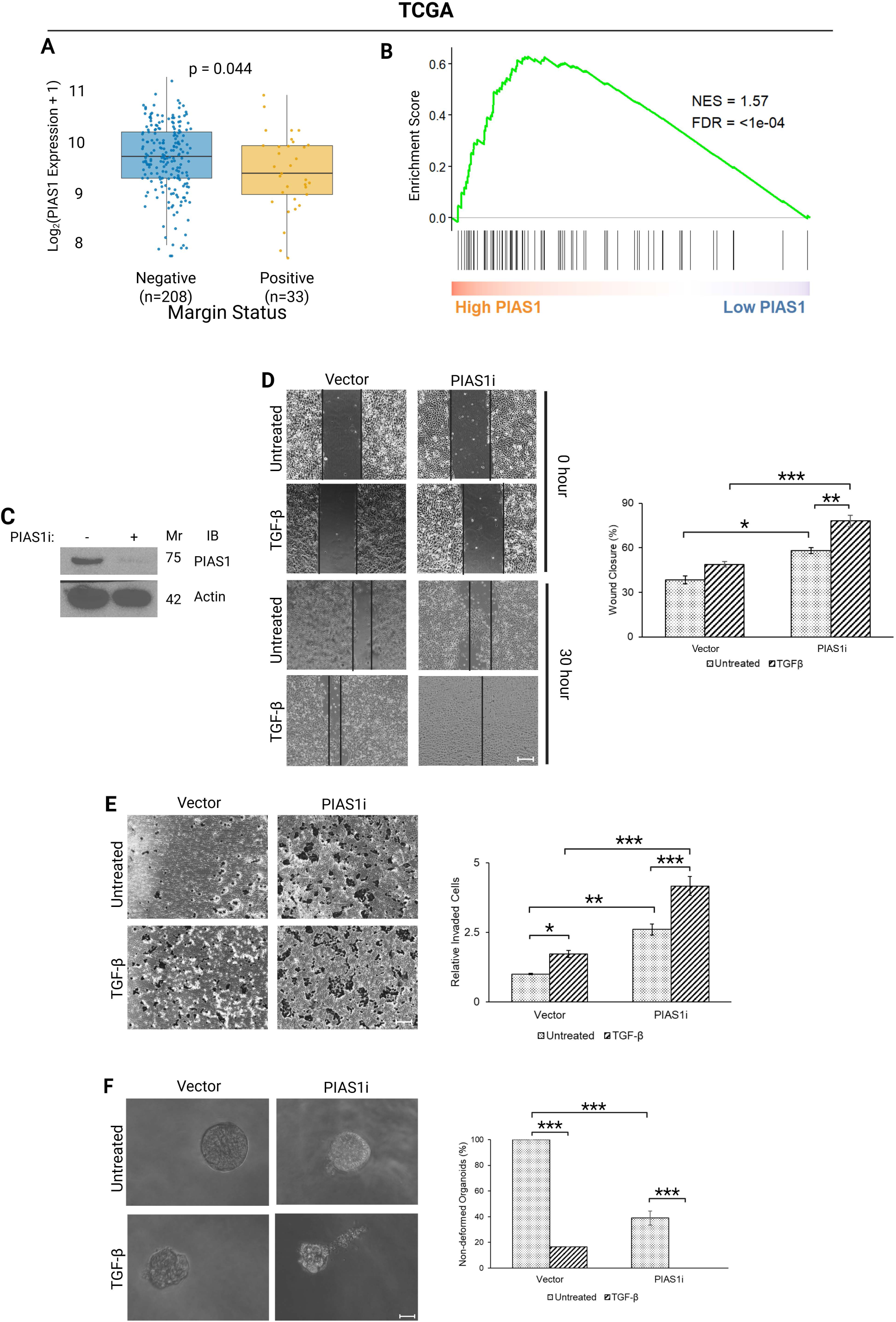
PIAS1 suppresses TGFβ-associated migration, invasion, and invasive growth of OSCC cells. (A) Box plot comparing PIAS1 expression in OSCC tumors with negative surgical margins (n=208) and positive surgical margins (n=33) in the TCGA cohort. (B) Gene set enrichment plots showing enrichment of TGFβ-induced EMT DOWN signature in High PIAS1 expressing tumors in the TCGA cohort. (C) Immunoblot showing PIAS1 expression in PIAS1-knockdown (PIAS1i) and control CAL33 cells, with actin used as a loading control. (D) Representative images from scratch assays performed on PIAS1i and control CAL33 cells, with or without TGFβ treatment, and corresponding quantification of wound closure at 30h. (E) Representative images from transwell invasion assays performed on PIAS1i and control CAL33 cells, with or without TGFβ treatment, and corresponding quantification of invaded cells at 24h. (F) Representative images of 8-day-old 3D organoids grown from PIAS1i and control CAL33 cells, with or without TGFβ treatment, and corresponding quantification of non-deformed organoids. *p<0.05, **p<0.01, ***p<0.001. Scale bars for scratch and transwell assays are 500 μm and scale bars for 3D culture are 50 μm.

To support the hypothesis that PIAS1 suppresses TGFβ-induced aggressive behaviour in OSCC, we used CAL33 cells, a well-characterized and widely used model system for functional studies, and for evaluating novel therapeutics and molecular biomarkers (48). PIAS1 was knocked down using PIAS1i (16, 17) (Fig. 2C). Control- or PIAS1i-vector-transfected CAL33 cells were left untreated or incubated with TGFβ and subjected to scratch and transwell invasion assays to compare cell migratory and invasion behaviour, respectively. As expected in the vector control, we found that CAL33 cells were able to migrate to close the scratch-generated gap, and TGFβ enhanced the migratory potential of these cells as indicated by the greater level of gap closure. Importantly, knockdown of endogenous PIAS1 promoted gap closure in both untreated and TGFβ treatment conditions (Fig. 2D). Similarly, in a transwell invasion assay, PIAS1i increased cell invasion even in the absence of TGFβ (Fig. 2E). Three-dimensional culturing of epithelial cells in Matrigel^TM^ provides an effective platform for modeling complex cell-cell interactions and promoting the formation of organoid-like structure. Compared with conventional two-dimensional culture systems, organoid cultures more faithfully recapitulate *in vivo* conditions, particularly in the context of TGFβ signaling and PIAS1 function (16–18). We therefore asked whether CAL33 could establish three-dimensional cultures and if so, what the role of PIAS1 is in this model. We found that isolated single CAL33 cells cultured in Matrigel^TM^ established spherical structures or organoids. Consistent with our migration and invasion findings, enhanced TGFβ-mediated signaling led to disrupted/invasive behaviour of these organoids. Interestingly, reduction of endogenous PIAS1 by knockdown disrupted organoid integrity, leading to invasive phenotypes even without TGFβ treatment (Fig. 2F). Collectively, these findings indicate that PIAS1 suppresses invasive and migratory potential of OSCC cells and helps maintain their overall epithelial organization.

We next investigated whether the effects of PIAS1 on OSCC cell migration and invasion depend on its SUMO E3 ligase activity. CAL33 cells were transfected with either wild-type PIAS1 (PIAS1 WT), a SUMO E3 ligase-inactive mutant PIAS1 CS (in which the catalytic cysteine at the 350th position is replaced by a serine (16)), or the appropriate empty plasmid (Suppl. Fig. 1B). TGFβ-induced scratch closure was significantly suppressed in PIAS1 WT-expressing cells, whereas PIAS1 CS expression promoted closure even without TGFβ (Suppl. Fig. 1C). Similarly, in transwell invasion assays, PIAS1 WT suppressed and PIAS1 CS promoted invasion (Suppl. Fig. 1D). In 3D culture, PIAS1 WT maintained organoid integrity, whereas PIAS1 CS promoted invasive growth (Suppl. Fig. 1E). These results suggest that PIAS1 suppresses OSCC cell invasiveness through its SUMO E3 ligase activity.

### Endogenous PIAS1 is inversely associated with gene signatures that predict poor OSCC outcomes

We then evaluated the downstream effects of PIAS1 expression in OSCC. We used control and PIAS1i expressing CAL33 cells in triplicate to perform RNA-seq (Fig. 3A) and identified 617 deregulated genes (≥1.5-fold change, p < 0.05) in PIAS1-depleted cells (Fig. 3B). Pathway analysis revealed upregulation of TGFβ signaling, cell cycle progression, and mitotic signals (Fig. 3C) that have been shown to be associated with tumor progression. To understand the functional relevance of these deregulated genes, we examined the spatial expression patterns of the upregulated and downregulated gene signatures separately in a cohort of 12 spatially profiled OSCC patient samples, in which we had previously identified tumor core, transitory and leading edge-associated expression programs (Fig. 3D, (42)). In aggregate UMAP projections of cancer cell-containing spatial spots from these 12 OSCC samples, genes upregulated following PIAS1 depletion were significantly enriched in the LE (Figure 3E), whereas the downregulated genes were enriched in the TC (Fig. 3F). These signatures were largely consistent across individual samples as well (Suppl. Fig. 2). Together, these findings suggest that PIAS1 regulates a transcriptional program linked to the aggressive behaviour of OSCC cells.

**Figure 3:**
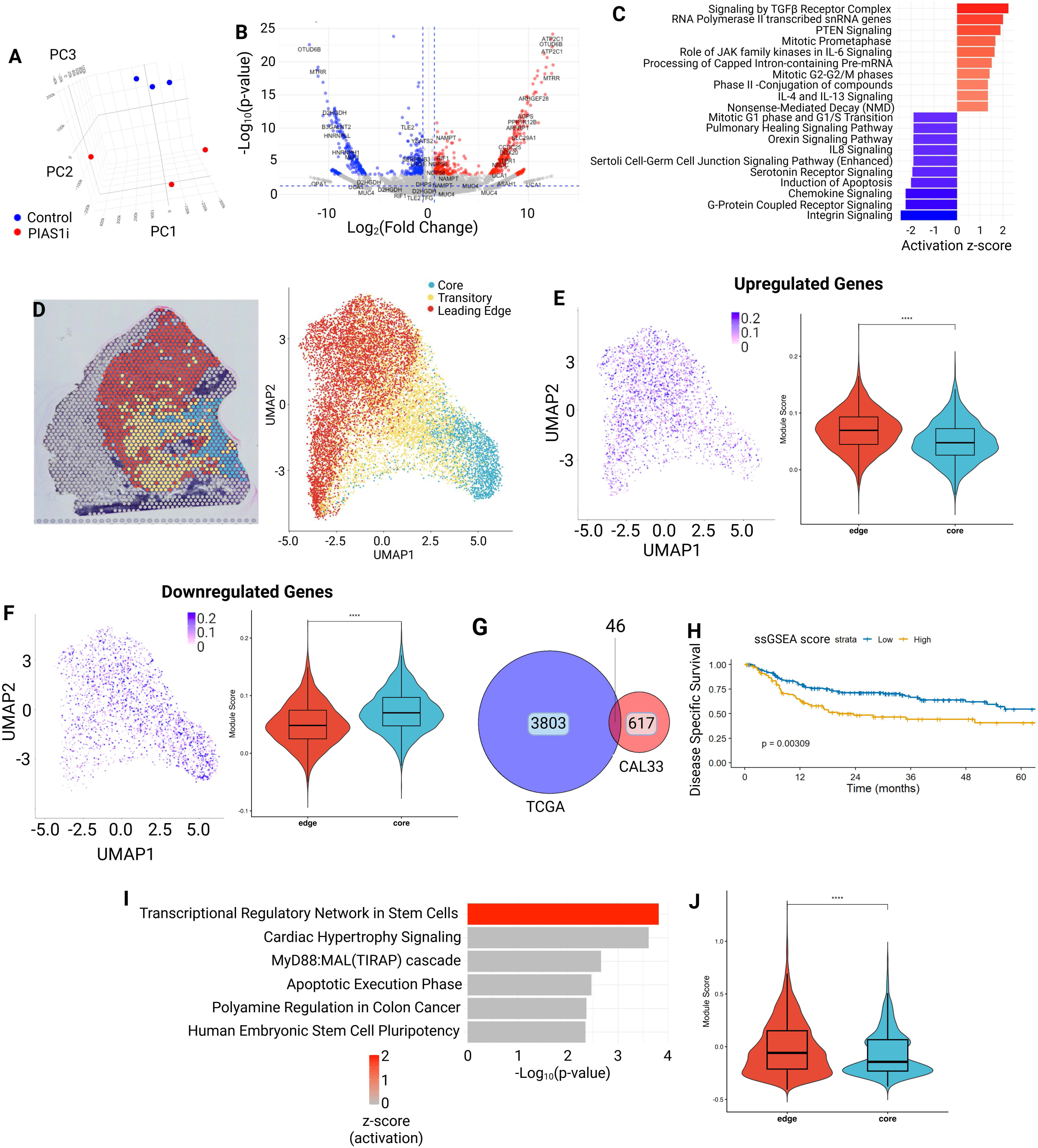
PIAS1 depletion identifies transcriptional programs associated with aggressive spatial tumor regions and poor OSCC outcomes. (A) 3D PCA plot of RNA-seq profiles from PIAS1-knockdown (PIAS1i) and control CAL33 cells used for differential gene expression analysis. (B) Volcano plot showing differentially expressed genes in PIAS1i cells compared with control cells, with selected highly deregulated genes labeled. (C) Waterfall plot showing the top upregulated and downregulated pathways identified by Ingenuity Pathway Analysis (IPA) of genes deregulated following PIAS1 knockdown. (D) Pathologist and gene signature based designation of tumor core, transitory, and leading-edge regions in one representative sample on the left and the UMAP projection of these regions from spatial transcriptomics data from all 12 samples. (E) UMAP visualization and violin plot showing module scores for genes upregulated following PIAS1 knockdown across leading edge and tumor core regions. (F) UMAP visualization and violin plot showing module scores for genes downregulated following PIAS1 knockdown across leading edge and tumor core regions. (G) Venn diagram showing overlap between genes deregulated in PIAS1-knockdown CAL33 cells and genes deregulated between PIAS1-low and PIAS1-high tumors in the TCGA OSCC cohort. (H) Kaplan-Meier curve for disease-specific survival (DSS) based on fold-change weighted ssGSEA scores of the common deregulated gene set. (I) Pathway analysis of the overlapping gene set showing enrichment of stemness-associated transcriptional regulatory programs. (J) Violin plot showing module scores for the four genes contributing to the top stemness-associated pathway across leading edge and tumor core regions in spatial transcriptomics data. ****p<0.0001.

To refine this gene signature to identify targetable molecules, we analyzed the TCGA OSCC cohort by stratifying tumors into tertiles based on PIAS1 expression. Specifically, we performed differential gene expression analysis between the bottom tertile with the lowest PIAS1 expression (n=92) and the top tertile with the highest expression (n=92) (Suppl. Fig. 3A), which identified 3803 deregulated genes (Suppl. Fig. 3B). Overlap of this gene set with the deregulated gene identified in CAL33 cells following PIAS1 inhibition, as described in Figure 3, revealed 46 common genes (Fig. 3G, Suppl. Table 3). We then performed ssGSEA on the 24 commonly upregulated and the 22 commonly downregulated genes. The gene set was significantly associated with worse DSS (Fig. 3H) and PFI in the TCGA OSCC cohort (Suppl. Fig. 3C). In contrast, the downregulated gene set was significantly associated with improved PFI (Suppl. Fig 3D). OS showed a similar trend for both signatures, although these associations did not reach statistical significance (data not shown). Pathway analysis identified only one significantly activated pathway (Figure 3I), which is linked to increased transcriptional activity of stemness-related programs. Four genes upregulated upon PIAS1 reduction (RIF1, INHBA, REST, and IL6ST) were found to drive this pathway. Notably, cancer cell stemness has been shown to be promoted by EMT across multiple tumor types, including OSCC (7, 49). Consistent with this, ssGSEA analysis of the four gene signature demonstrated significant enrichment in the LE as compared to the TC (Fig. 3J, Suppl. Fig. 3E), a pattern that was generally preserved across individual samples (Suppl. Fig. 3F). Collectively, these findings indicate that this stemness-associated pathway and its linked four-gene signature are associated with aggressive behaviour in OSCC tumors.

### INHBA marks invasive OSCC states and defines a reciprocal INHBA-PIAS1 phenotype

To further define the individual genes emerging from the PIAS1-regulated stemness-associated signature, we examined each candidate across independent OSCC and HNSCC cohorts. Among these, INHBA emerged as a particularly strong candidate because it showed consistent tumor-associated upregulation, association with nodal metastasis, inverse relationship with PIAS1, and spatial enrichment within invasive tumor regions. In TCGA OSCC, INHBA expression was significantly higher in tumors compared with normal oral cavity squamous epithelium and was further increased in node-positive tumors compared with node-negative tumors (Fig. 4A, B). These findings were reproduced in the Calgary RNA-seq cohort, where INHBA was similarly elevated in tumors relative to matched normal tissues and in tumors from patients with nodal metastasis (Suppl. Fig. 4A, B). Consistent with the hypothesis that INHBA is linked to reduced PIAS1 activity, INHBA expression showed an inverse correlation with PIAS1 expression in TCGA tumors (Suppl. Fig. 4C).

**Figure 4:**
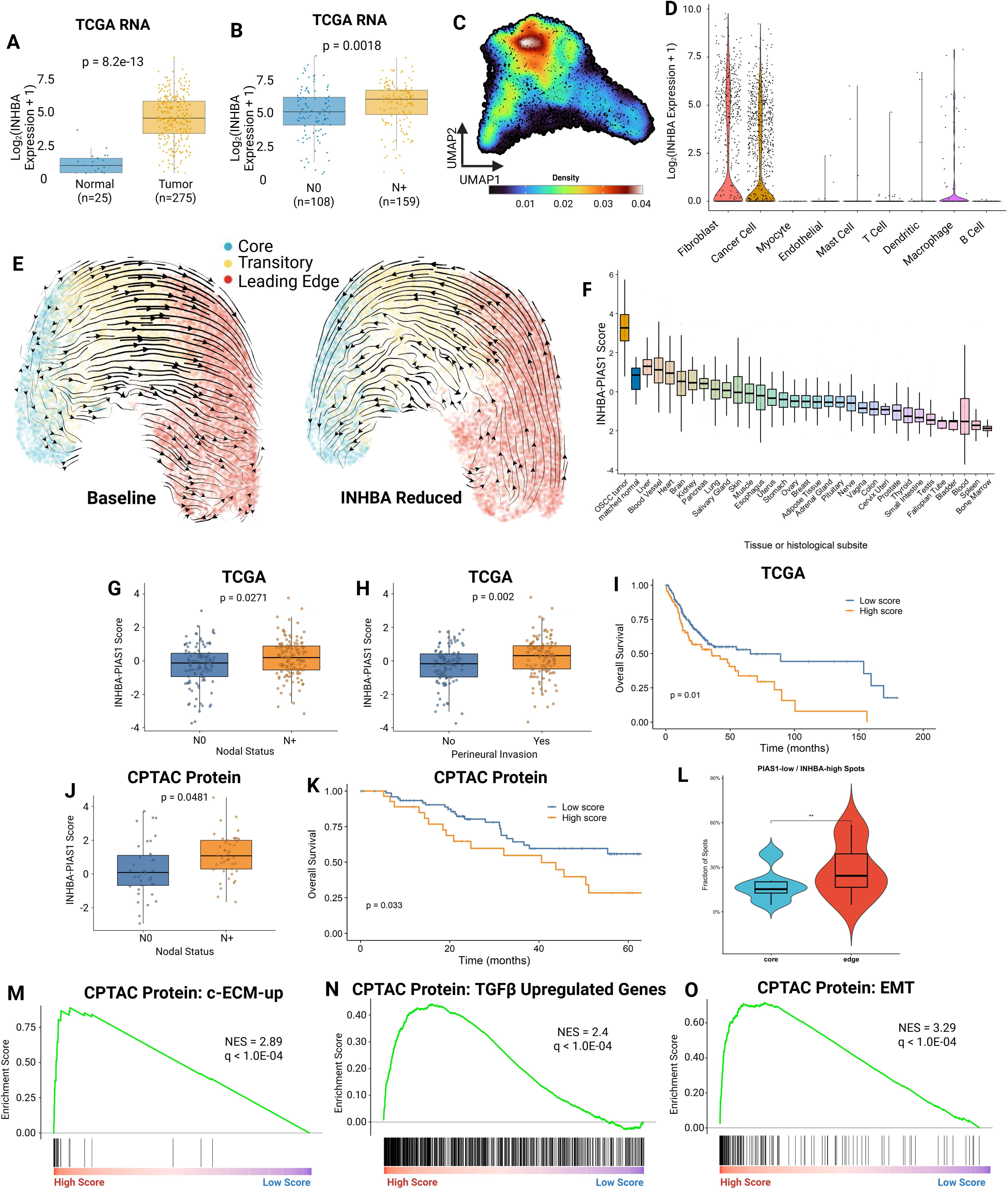
INHBA marks invasive OSCC states and defines a reciprocal INHBA-PIAS1 phenotype. (A) Box plot showing INHBA expression in OSCC tumor samples compared to normal OCSE in the TCGA cohort. (B) Box plot showing INHBA expression in node-negative (N0) and node-positive (N1+) primary OSCC tumors in the TCGA cohort. (C) Spatial density plot showing INHBA expression across spatial transcriptomics-derived tumor regions. (D) Violin plot showing INHBA expression across cell types in single-cell RNA-seq analysis of the OSCC tumor microenvironment. (E) UMAP of spatially deconvolved cancer-containing spots with overlaid RNA velocity streams, colored according to tumor core, transitory, and leading-edge annotations under baseline and INHBA-reduced conditions. (F) Box plot showing the INHBA-PIAS1 score across OSCC and normal body subsites. The INHBA-PIAS1 score was calculated so that higher values reflect relatively higher INHBA and lower PIAS1 abundance. (G) Box plot comparing the INHBA-PIAS1 score in node-negative (N0) and node-positive (N+) TCGA OSCC tumors. (H) Box plot comparing the INHBA-PIAS1 score in TCGA OSCC tumors with or without perineural invasion. (I) Kaplan-Meier curve for OS based on high and low INHBA-PIAS1 score groups in the TCGA OSCC cohort. (J) Box plot comparing the INHBA-PIAS1 protein score in node-negative (N0) and node-positive (N+) tumors in the CPTAC cohort. (K) Kaplan-Meier curve for OS based on high and low INHBA-PIAS1 protein score groups in the CPTAC cohort. (L) Box plot showing the fraction of spatial spots with high INHBA-PIAS1 scores across tumor core and leading-edge regions, with matched samples connected by lines. (M-O) Gene set enrichment plots showing enrichment of C-ECM-up, TGFβ-upregulated genes, and EMT programs in high versus low INHBA-PIAS1 score tumors using CPTAC protein data.

We next asked whether this relationship was preserved at the protein level. In the CPTAC HNSCC cohort, INHBA protein abundance was significantly higher in tumors than in normal tissues and was inversely correlated with PIAS1 protein abundance (Suppl. Fig. 4D, E). Higher INHBA protein abundance was also associated with worse overall survival in CPTAC, supporting the clinical relevance of INHBA at the protein level (Suppl. Fig. 4F). Together, these transcriptomic and proteomic analyses indicate that INHBA is upregulated in OSCC/HNSCC tumors, is associated with metastatic progression, and is negatively linked to PIAS1 abundance across independent datasets.

Given the association of INHBA with aggressive clinicopathologic features, we next examined its spatial and cellular distribution in the OSCC tumor microenvironment. In our spatial transcriptomics cohort, INHBA expression was enriched toward the leading edge, a compartment previously linked to invasive tumor behavior and poor clinical outcome (Fig. 4C). Single-cell RNA-seq analysis of primary HNSCC tumors (23) showed that INHBA was expressed predominantly in fibroblasts and malignant epithelial cells, with relatively limited expression in most immune and endothelial compartments (Fig. 4D). A greater proportion of cancer-associated fibroblasts and malignant epithelial cells expressed INHBA compared with normal fibroblasts and normal epithelial cells, respectively (Suppl. Fig. 4G). Thus, INHBA appears to arise from both malignant and stromal compartments, positioning it as a candidate mediator of tumor-intrinsic invasion and tumor-stromal remodeling. This pattern was notable given our previous single-cell analysis showing that PIAS1-expressing CAFs and cancer cells were significantly reduced compared with their corresponding normal fibroblast and epithelial compartments (50).

We then asked whether INHBA contributes to spatial tumor-state transitions. RNA velocity leverages the ratio of spliced to unspliced mRNA to predict the short-term future states of individual cells (51). When aggregated across cells, these predictions can reveal developmental trajectories of cancer cells and help identify genes that drive state transitions (52). RNA velocity analysis of spatially profiled cancer-containing spots showed a trajectory from the tumor core toward the leading edge under baseline conditions (Fig. 4E, Baseline; Suppl. Fig. 4H). In silico reduction of INHBA using Dynamo redirected the inferred vector field away from the leading edge and toward the tumor core (Fig. 4E, INHBA reduced; Suppl. Fig. 4H). Although this perturbation analysis is computational and requires experimental validation, it supports the possibility that INHBA helps maintain the transcriptional dynamics of the invasive front.

Because the preceding analyses suggested a reciprocal relationship between increased INHBA and reduced PIAS1, we generated an INHBA-PIAS1 score, in which higher values represent relatively higher INHBA abundance together with relatively lower PIAS1 abundance. This score was designed to capture the biological state suggested by our functional and patient-cohort analyses, namely reduced PIAS1-associated restraint coupled to increased INHBA-associated invasive signaling. Across TCGA and GTEx tissue groups, the INHBA-PIAS1 score was elevated in OSCC compared with matched normal tissue and most normal tissue types (Fig. 4F). In contrast, INHBA alone showed overlap with selected normal tissues, including vascular-rich tissues, while PIAS1 alone did not show the same tumor-selective pattern (Suppl. Fig. 5A, B). Consistent with the construction of the composite score, tumors in the high INHBA-PIAS1 score group showed significantly lower PIAS1 expression and significantly higher INHBA expression than tumors in the low-score group in the TCGA OSCC cohort (Suppl. Fig. 5C, D). These findings suggest that the composite score may better identify an OSCC-enriched biological state than INHBA expression alone, while also indicating that therapeutic strategies directed at INHBA would require careful assessment of normal-tissue expression.

We next assessed whether the INHBA-PIAS1 score was associated with clinical aggressiveness. In TCGA OSCC, the score was significantly higher in node-positive tumors than in node-negative tumors and was also higher in tumors with perineural invasion (Fig. 4G, H). Patients with high INHBA-PIAS1 scores had worse overall survival compared with those with low scores in TCGA (Fig. 4I). The same direction of association was observed in CPTAC, where high INHBA-PIAS1 protein scores were associated with nodal metastasis and worse overall survival (Fig. 4J, K). RNA-level CPTAC analyses further supported the association of high INHBA-PIAS1 score with nodal involvement, advanced stage, and a trend toward reduced survival (Suppl. Fig. 5E-G). In our spatial transcriptomics cohort, the fraction of high INHBA-PIAS1 spots was increased in LE compared with TC regions (Fig. 4L). Collectively, these analyses show that the INHBA-high/PIAS1-low state identifies a clinically aggressive and spatially invasive OSCC phenotype.

Finally, we examined whether the INHBA-PIAS1 score was associated with biological pathways expected from the functional PIAS1 and INHBA data. In CPTAC proteomic data, tumors with high INHBA-PIAS1 scores showed enrichment of cancer-associated extracellular matrix remodeling, TGFβ-upregulated genes, and EMT-related programs (Fig. 4M-O). Similar enrichment patterns were observed at the RNA level in TCGA and CPTAC datasets (Suppl. Fig. 5H-L). These findings connect the composite score to the same biological processes implicated by PIAS1 loss and INHBA upregulation, supporting a model in which the INHBA-high/PIAS1-low state reflects TGFβ-linked EMT, matrix remodeling, and invasive tumor behavior.

### The INHBA-PIAS1 score is associated with immunotherapy resistance and fibroblast-dominant immune suppression

Because extracellular matrix remodeling and TGFβ-associated stromal programs have been linked to immune exclusion and reduced response to immune checkpoint blockade, we next asked whether the INHBA-PIAS1 score was associated with immunotherapy outcome. Across seven immune checkpoint blockade-treated patient cohorts, non-responders had significantly higher INHBA-PIAS1 scores than responders (Fig. 5A). This suggested that the INHBA-high/PIAS1-low phenotype may not only mark invasive disease but may also identify tumors with reduced sensitivity to immune checkpoint therapy.

**Figure 5:**
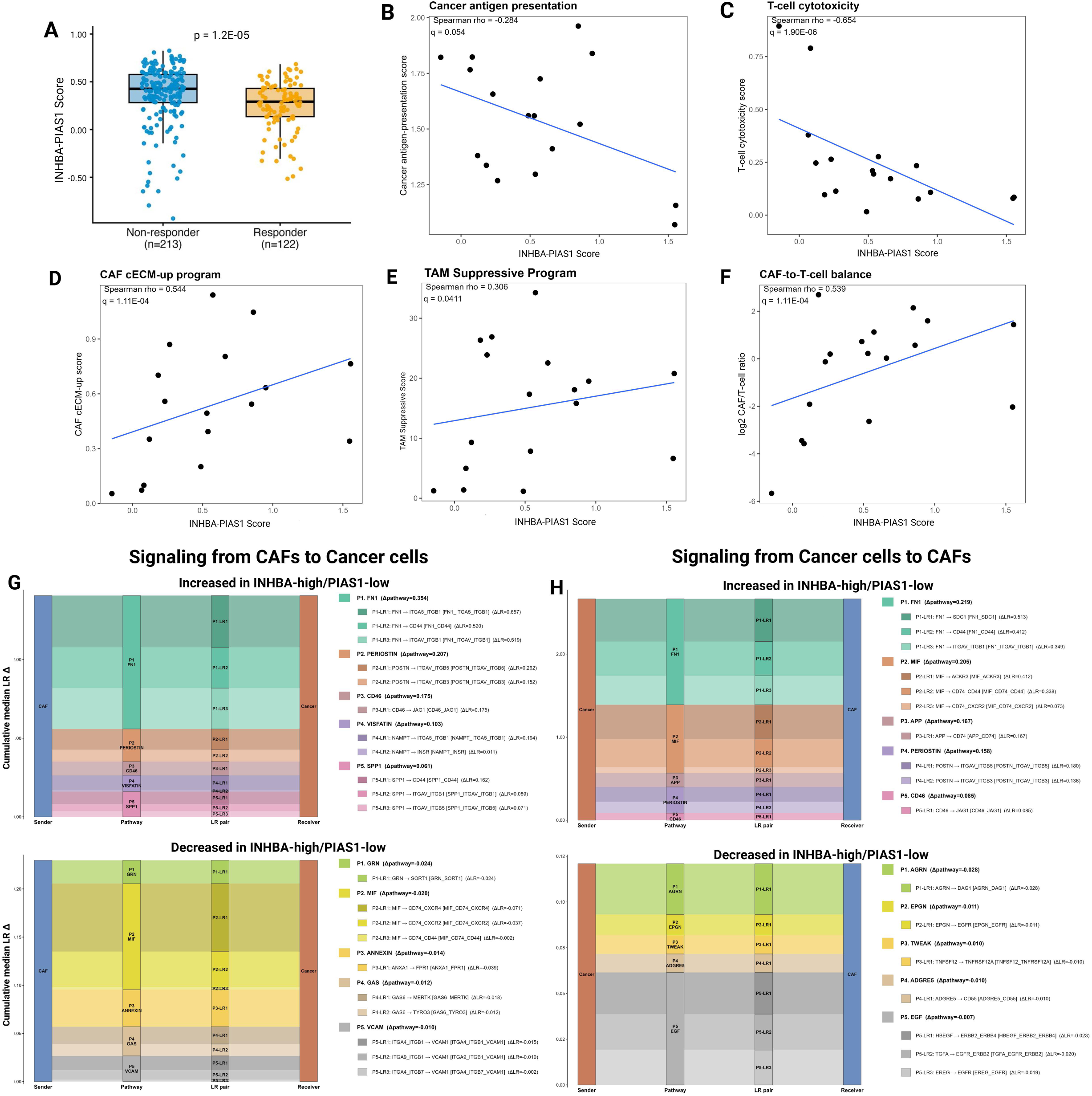
The INHBA-PIAS1 score is associated with immunotherapy resistance and a matrix-centered, immune-suppressive CAF-cancer communication circuit. (A) Box plot showing INHBA-PIAS1 scores in immunotherapy non-responders (n=213) and responders (n=122) pooled across seven immune checkpoint blockade-treated cohorts. (B to F) Scatter plots showing Spearman correlations between the INHBA-PIAS1 score and (B) cancer cell antigen-presentation score, (C) T-cell cytotoxicity score, (D) CAF c-ECM-up program score, (E) TAM suppressive program score, and (F) log2 CAF-to-T-cell ratio, with Spearman rho and q-values indicated. (G, H) Differential ligand–receptor communication between CAF and cancer-cell compartments in the spatial data, using ligand–receptor pairs curated in CellChatDB, comparing INHBA-high/PIAS1-low with INHBA-low/PIAS1-high states, shown separately for (G) signaling from CAFs to cancer cells and (H) signaling from cancer cells to CAFs. Each alluvial plot links sender, signaling pathway, ligand-receptor pair, and receiver. The top plot in each panel shows interactions increased and the bottom plot shows interactions decreased in the INHBA-high/PIAS1-low state. The y-axis (cumulative median LR Δ) and bar height represent the difference in communication strength between states. Δpathway denotes the aggregated pathway-level change and ΔLR the change for each ligand-receptor pair, with positive values indicating stronger signaling in INHBA-high/PIAS1-low tumors.

To define the tumor microenvironment associated with this state, we first correlated the INHBA-PIAS1 score with cell-type and functional programs in spatial transcriptomic data. ECM-CAF and myofibroblasts were enriched in the tumor microenvironment of high INHBA-PIAS1 score containing tumor samples, suggesting an immune excluded microenvironment (Suppl. Fig. 5M). In single-cell RNA-seq data, tumors with high INHBA-PIAS1 scores showed lower cancer cell antigen presentation and T-cell cytotoxicity scores (Fig. 5B, C). In contrast, high-score tumors showed higher cancer-associated fibroblast C-ECM-up scores, increased suppressive tumor-associated macrophage features, and increased CAF-to-T-cell ratios (Fig. 5D, E, F). These results indicate that the INHBA-high/PIAS1-low phenotype is associated with a stromal-rich and immune-suppressed microenvironment.

We next evaluated whether these relationships were also evident across spatially resolved pathway-level signatures. The INHBA-PIAS1 score showed positive correlations with hypoxia, myCAF, TGFβ signaling, CAF/ECM remodeling, and EMT signatures, while showing negative correlations with MHC-II antigen presentation, TIL/T-cell activation, and CD8 cytotoxicity-associated signatures (Suppl. Fig. 5N). Thus, the score is linked to a coordinated microenvironmental state characterized by stromal remodeling, TGFβ-associated invasion, and impaired anti-tumor immune activity.

Finally, we used CellChat to ask whether the INHBA-high/PIAS1-low state reshapes signaling between CAFs and malignant cells, and in which direction. In the CAF-to-cancer direction, the dominant gains in INHBA-high/PIAS1-low tumors were matrix and adhesion pathways, led by FN1 (Δpathway = 0.354) and followed by Periostin, CD46, Visfatin (NAMPT), and SPP1, acting largely through integrin and CD44 receptors on cancer cells, for example FN1 to ITGA5_ITGB1 (ΔLR = 0.657) and to CD44 (ΔLR = 0.520), and SPP1 to CD44 (ΔLR = 0.162) (Fig. 5G, top). The reciprocally reduced interactions in this direction, involving progranulin (GRN), MIF, Annexin, GAS, and VCAM, were all of small magnitude (|Δpathway| ≤ 0.024) (Fig. 5G, bottom). In the cancer-to-CAF direction, INHBA-high/PIAS1-low tumors gained FN1 (Δpathway = 0.219) together with MIF (Δpathway = 0.205), APP, Periostin, and CD46 signaling, including MIF to CD74-containing receptor complexes and APP to CD74 (Fig. 5H, top), while the decreased pathways, comprising EGF-family ligands (HBEGF, TGFA, EREG), TWEAK, AGRN, EPGN, and ADGRE5, were again low in magnitude (|Δpathway| ≤ 0.028) (Fig. 5H, bottom). The increased communication was therefore bidirectional and carried by shared ECM and adhesion pathways (FN1, Periostin, CD46) plus direction-specific programs, namely SPP1 and Visfatin from CAFs and MIF and APP from cancer cells. These patterns indicate that the INHBA-high/PIAS1-low state reinforces a matrix- and adhesion-centered tumor-stromal communication circuit rather than a purely tumor-cell-autonomous phenotype.

Collectively, these analyses extend the Figure 4 findings by showing that the INHBA-high/PIAS1-low state is associated with immunotherapy non-response, CAF enrichment, suppressive macrophage features, reduced T-cell cytotoxicity, and strengthened, matrix-centered bidirectional CAF-cancer communication. These data support a model in which loss of PIAS1 and gain of INHBA converge on an invasive, ECM-remodeling, and immune-suppressed OSCC state with potential implications for patient stratification and therapeutic targeting.

## Discussion

In this study, we identify PIAS1 as a tumor-suppressive regulator of OSCC progression that acts, at least in part, by suppressing TGFβ-linked EMT, invasion, and extracellular matrix remodeling. Across independent patient cohorts, PIAS1 mRNA and protein abundance were reduced in tumors compared with normal tissues, and higher PIAS1 protein abundance was associated with improved patient outcome. Functional assays further showed that PIAS1 suppresses OSCC cell migration, invasion, and invasive growth in three-dimensional culture in a SUMO E3 ligase-dependent manner. These findings support a model in which reduced PIAS1 activity removes a regulatory brake on pro-invasive transcriptional programs in OSCC.

The cancer-related function of PIAS1 is highly context dependent. Previous studies have reported both tumor-promoting and tumor-suppressive roles for PIAS1, likely reflecting differences in tumor type, substrate availability, signaling context, and the specific cellular processes under PIAS1 control (14, 16, 17, 53–56). In breast epithelial and breast cancer models, PIAS1 has been shown to suppress TGFβ-induced EMT through SUMO-dependent mechanisms involving regulators such as SnoN (Ski-related novel protein N) (15–18). Our findings extend this concept to OSCC and suggest that PIAS1-mediated sumoylation may similarly oppose TGFβ-driven invasive behavior in oral cancer cells. Future work should determine whether canonical PIAS1 substrates, including SnoN or other TGFβ pathway regulators, mediate the anti-invasive effects of PIAS1 in OSCC.

A major finding from this study is the identification of INHBA as a clinically and biologically relevant downstream component of the PIAS1-associated invasive program. INHBA encodes the inhibin beta A subunit and has been linked to tumor progression, EMT, stromal activation, immune modulation, and treatment resistance in several cancers (57–61). In HNSCC, elevated INHBA expression correlates with lymph node metastasis, chemoresistance, and poor survival outcomes (61, 62). In our analyses, INHBA was upregulated in OSCC tumors compared with normal tissues, increased in node-positive disease, inversely associated with PIAS1 expression, and associated with worse survival at the protein level. Spatial transcriptomics further showed enrichment of INHBA at the leading edge, while single-cell analysis identified malignant epithelial cells and cancer-associated fibroblasts as major INHBA-expressing compartments. These findings suggest that INHBA may contribute to OSCC progression through both tumor-cell-intrinsic and stromal-associated mechanisms.

The spatial localization of INHBA is particularly relevant because the leading edge represents a biologically aggressive tumor compartment enriched for invasive and therapy-resistant programs (42). RNA velocity analysis supported a baseline transition from tumor core toward leading-edge states, and in silico INHBA reduction redirected this inferred trajectory away from the leading edge and toward the tumor core. Although this perturbation approach is computational and should not be interpreted as direct experimental proof, it provides a mechanistic rationale for studying INHBA as a regulator of invasive tumor-state maintenance. Functional perturbation of INHBA in OSCC models, ideally in tumor-CAF co-culture, organoid, and in vivo systems, will be required to validate this prediction.

An important translational point from our analysis is that INHBA expression alone may not fully define a tumor-selective therapeutic context because INHBA expression overlaps with selected normal tissues. To address this, we developed a reciprocal INHBA-PIAS1 score that captures the combined state of high INHBA and low PIAS1. This composite score was enriched in OSCC, associated with nodal metastasis, perineural invasion, poor survival, leading-edge localization, and enrichment of EMT, TGFβ, and C-ECM-up programs. Thus, the score appears to capture a broader biological state than either marker alone, linking PIAS1 loss and INHBA gain to a coherent invasive phenotype. This may be useful for patient stratification, particularly if INHBA-directed therapies or TGFβ/ECM-targeted combination strategies are explored in OSCC.

The immune findings further extend this model. Previous research has established that INHBA influences the immune landscape by promoting an immunosuppressive environment, as observed in different cancer types (60, 63, 64). In our analyses, high INHBA-PIAS1 scores were associated with immunotherapy non-response, ECM-CAF and myofibroblast programs, suppressive macrophage features, reduced T-cell cytotoxicity, and increased CAF-to-T-cell ratios. Spatial signature analyses similarly linked the score to hypoxia, CAF/ECM remodeling, TGFβ signaling, and EMT, while showing negative associations with antigen presentation and cytotoxic immune programs. Ligand–receptor analysis of the spatial transcriptomic data suggested that high-score tumors have reinforced bidirectional communication between CAFs and malignant cells, dominated by ECM and adhesion signaling. Fibronectin (FN1) was the strongest gain in both directions, accompanied by Periostin and CD46 in both directions, by SPP1 and Visfatin (NAMPT) from CAFs to cancer cells, and by MIF and APP from cancer cells to CAFs. The reciprocally decreased interactions were consistently of much smaller magnitude, so this state is better described as a gain of matrix- and adhesion-centered communication than as a wholesale rewiring. Several of these axes, in particular FN1-integrin, SPP1-CD44, Periostin-integrin, and MIF-CD74, are established contributors to invasion, ECM remodeling, and myeloid-mediated immune suppression (65–68), consistent with the invasive, immune-excluded phenotype associated with the INHBA-high/PIAS1-low state. These results support a model in which the INHBA-high/PIAS1-low state promotes not only invasion, but also stromal remodeling and immune suppression.

These observations are consistent with prior work linking TGFβ-associated extracellular matrix programs to immune exclusion and immunotherapy failure (69). In this context, INHBA may function as part of a broader stromal-immunosuppressive circuit rather than as an isolated tumor-intrinsic oncogene. This distinction is important therapeutically. Direct INHBA inhibition may be most effective in tumors with high INHBA-PIAS1 scores and evidence of CAF-rich, TGFβ-linked immune suppression. Alternatively, INHBA-directed therapy may need to be combined with immune checkpoint blockade, TGFβ pathway inhibition, macrophage-reprogramming strategies, or ECM-targeted approaches to overcome the multicellular architecture of the resistant tumor microenvironment.

This study has several limitations. First, although the association between PIAS1 loss, INHBA upregulation, and aggressive OSCC phenotypes is supported across multiple datasets, direct functional validation of INHBA as a mediator of PIAS1-loss phenotypes remains necessary. Second, the RNA velocity and Dynamo perturbation analyses provide hypothesis-generating evidence of INHBA-dependent spatial state changes, but these computational predictions require experimental validation. Third, the immunotherapy analyses integrate cohorts from different tumor types and treatment contexts, which supports generalizability but may introduce cohort-specific confounding. Finally, while the INHBA-PIAS1 score may improve biological specificity compared with INHBA alone, its clinical utility will require validation in independent OSCC cohorts with standardized treatment, spatial profiling, and outcome annotation.

In summary, our findings support a model in which PIAS1 inhibits TGFβ-linked invasive behavior in OSCC through SUMO E3 ligase-dependent mechanisms, while reduced PIAS1 is associated with increased INHBA and activation of an invasive stromal-immune program. The INHBA-high/PIAS1-low state identifies tumors with leading-edge enrichment, ECM remodeling, EMT activation, immune suppression, and poor clinical outcome (Graphical Abstract). These data nominate the PIAS1-INHBA axis as a biologically coherent and potentially targetable pathway in OSCC progression, while highlighting the need for functional INHBA inhibition studies and prospective validation of the INHBA-PIAS1 score.

## Supporting information

Full Blots

---

**Supplementary Figure 1:**
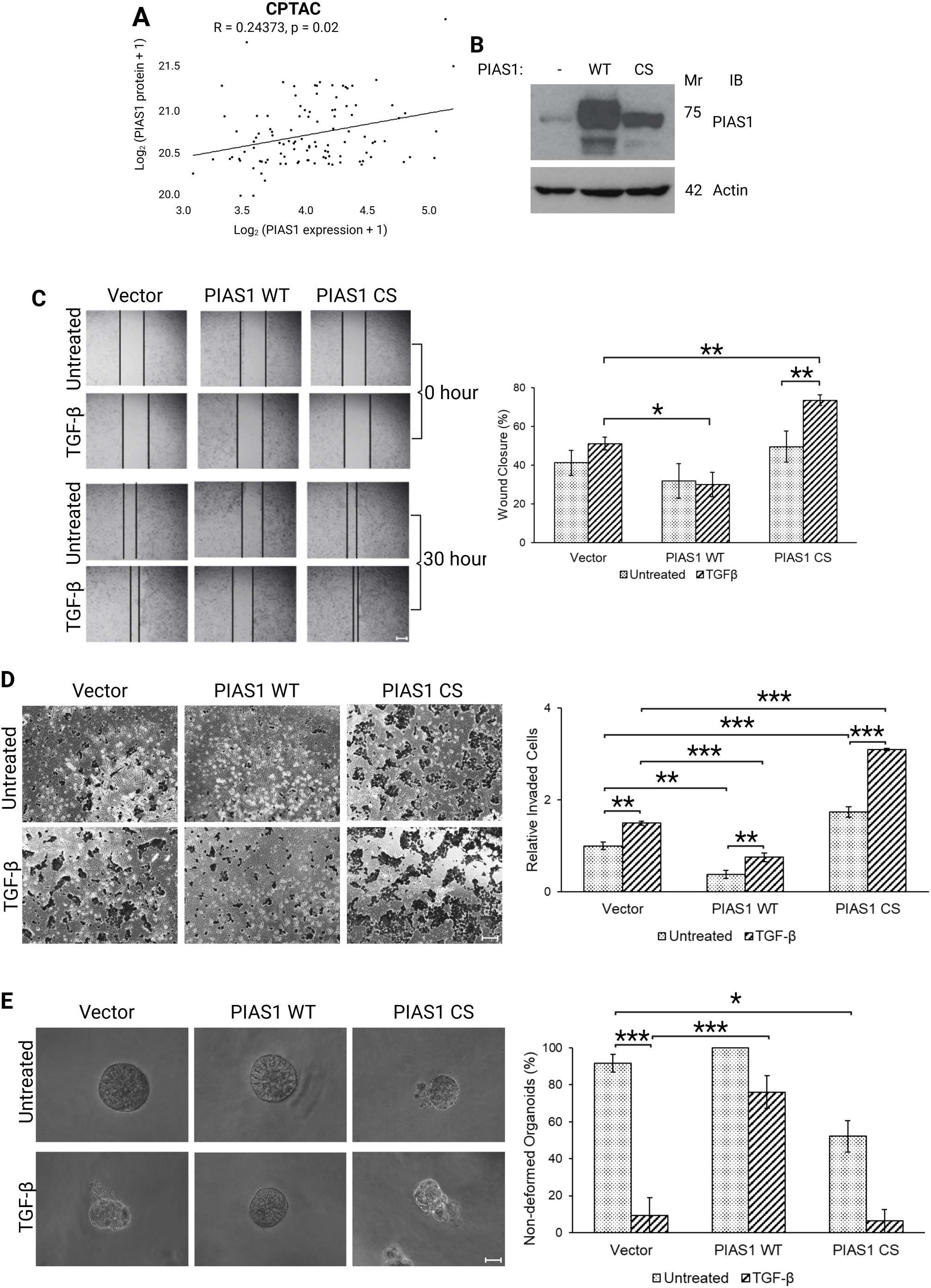
PIAS1 suppresses OSCC cell invasion through its SUMO E3 ligase activity. (A) Scatter plot showing Spearman correlation between PIAS1 mRNA expression and PIAS1 protein abundance in OSCC tumor samples from the CPTAC dataset. (B) Immunoblot showing PIAS1 expression in CAL33 cells transfected with wild-type PIAS1 (PIAS1 WT), SUMO E3 ligase-inactive PIAS1 (PIAS1 CS), or empty vector. (C) Representative images from scratch assays performed on PIAS1 WT, PIAS1 CS, and control CAL33 cells, with or without TGFβ treatment, and corresponding quantification of wound closure at 30h. (D) Representative images from transwell invasion assays performed on PIAS1 WT, PIAS1 CS, and control CAL33 cells, with or without TGFβ treatment, and corresponding quantification of invaded cells at 24h. (E) Representative images of 8-day-old 3D organoids grown from PIAS1 WT, PIAS1 CS, and control CAL33 cells, with or without TGFβ treatment, and corresponding quantification of non-deformed organoids. *p<0.05, **p<0.01, ***p<0.001.

**Supplementary Figure 2:**
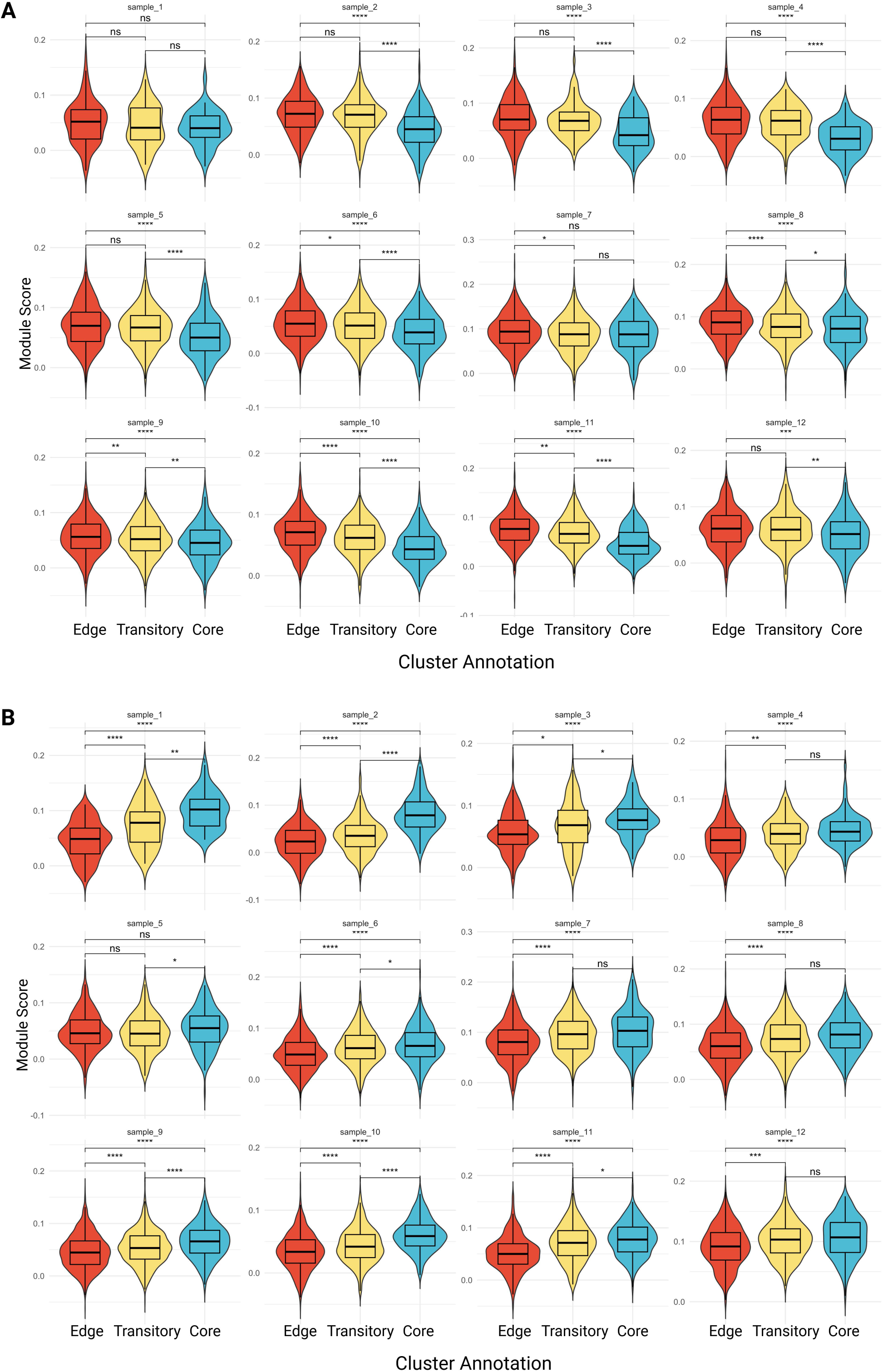
PIAS1-regulated gene signatures show spatial enrichment patterns across individual OSCC samples. (A) Sample-wise violin plots showing module scores for genes upregulated following PIAS1 knockdown across leading edge, transitory, and tumor core regions in spatial transcriptomics data. (B) Sample-wise violin plots showing module scores for genes downregulated following PIAS1 knockdown across leading edge, transitory, and tumor core regions in spatial transcriptomics data. ns = nonsignificant, *p<0.05, **p<0.01, ***p<0.001, ****p<0.0001.

**Supplementary Figure 3:**
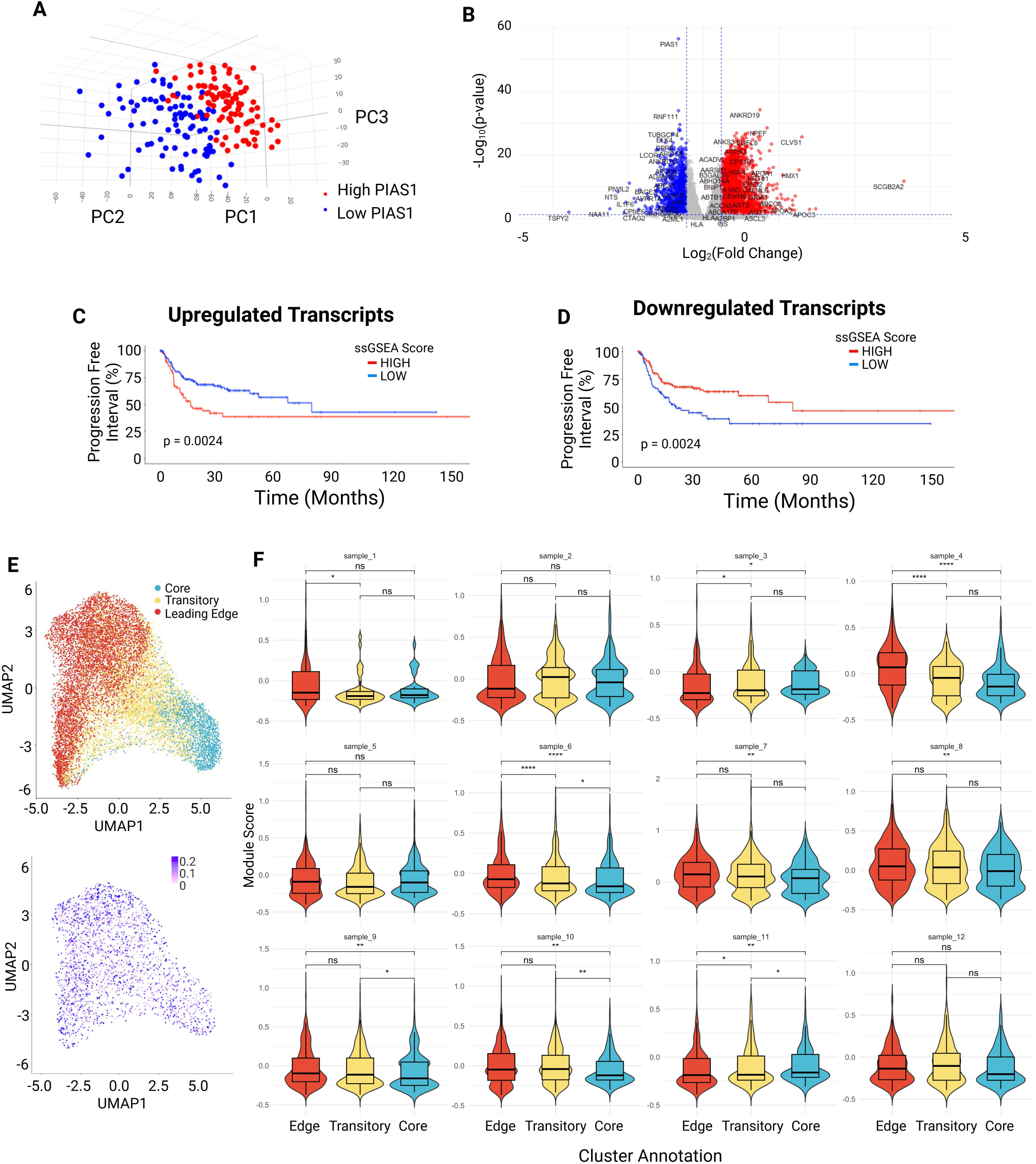
TCGA-derived PIAS1-associated transcriptional programs are linked to OSCC progression and spatial tumor states. (A) 3D PCA plot showing separation of TCGA OSCC tumors with high and low PIAS1 expression. (B) Volcano plot showing genes differentially expressed between PIAS1-low and PIAS1-high tumors in the TCGA OSCC cohort. (C) Kaplan-Meier curve for progression-free interval (PFI) based on ssGSEA scores of the commonly upregulated gene set. (D) Kaplan-Meier curve for PFI based on ssGSEA scores of the commonly downregulated gene set. (E) UMAP projection of spatial transcriptomics data showing tumor core, transitory, and leading-edge regions, with corresponding module score visualization for the four-gene stemness-associated signature. (F) Sample-wise analysis of module scores for the four-gene stemness-associated signature across leading edge, transitory, and tumor core regions in spatial transcriptomics data. ns = nonsignificant, *p<0.05, **p<0.01, ****p<0.0001.

**Supplementary Figure 4:**
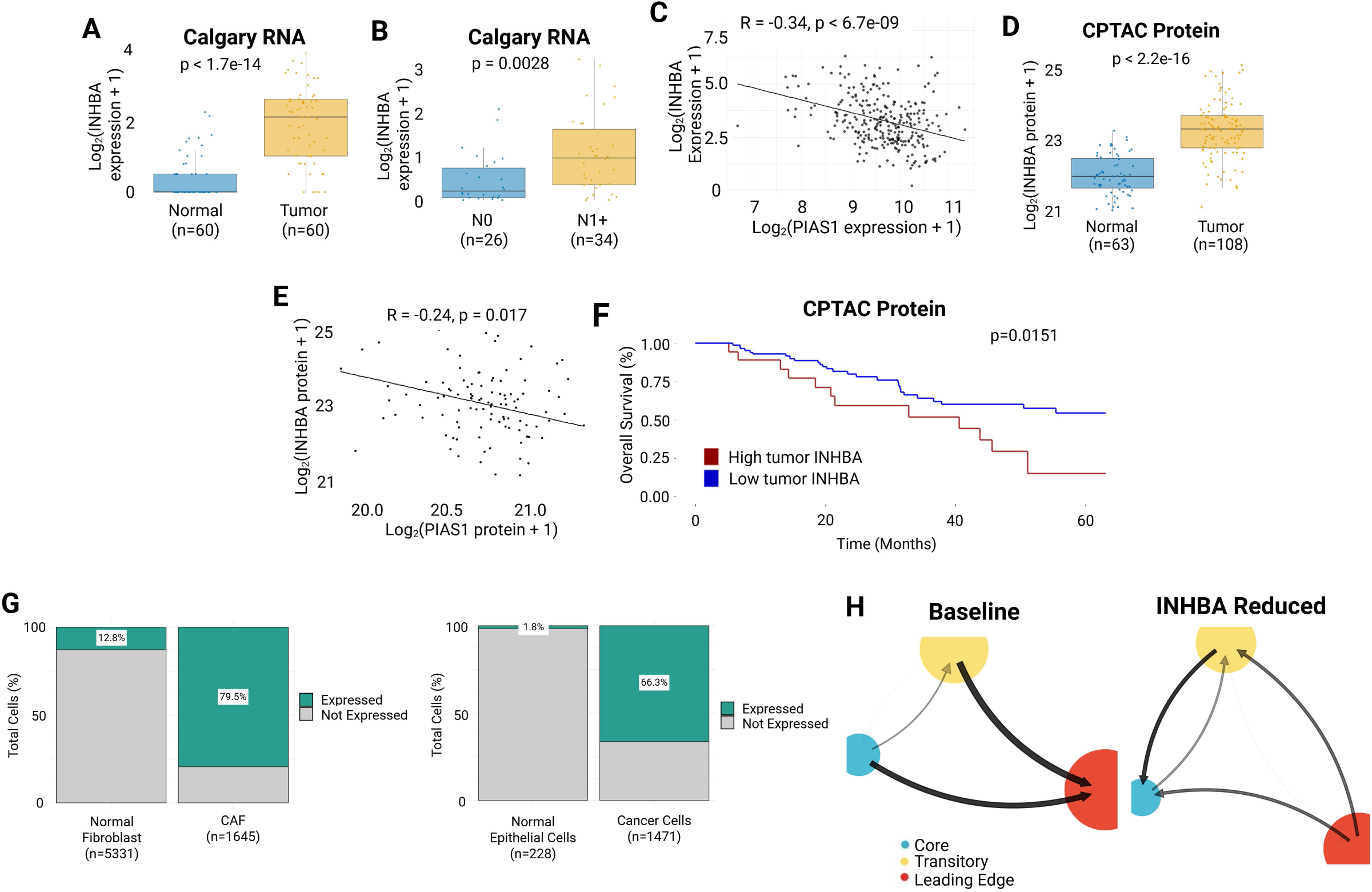
INHBA is upregulated in OSCC tumors, inversely associated with PIAS1, and enriched in malignant and stromal tumor compartments. (A) Box plot showing INHBA expression in matched OSCC tumor and normal samples in the Calgary OSCC cohort. (B) Box plot showing INHBA expression in node-negative (N0) and node-positive (N1+) primary tumors in the Calgary OSCC cohort. (C) Scatter plot showing Spearman correlation between INHBA and PIAS1 expression in tumor samples from the TCGA cohort. (D) Box plot showing INHBA protein abundance in tumor and normal samples in the CPTAC cohort. (E) Scatter plot showing Spearman correlation between INHBA and PIAS1 protein abundance in tumor samples from the CPTAC cohort. (F) Kaplan-Meier curve for OS based on INHBA protein abundance in patients from the CPTAC cohort, dichotomized into high and low tumor INHBA groups. (G) Stacked bar plots showing the proportion of INHBA-expressing cells in normal fibroblasts and CAFs, and in normal epithelial cells and malignant cancer cells, from single-cell RNA-seq analysis. (H) Cell fate transition probability state graph for tumor core, transitory, and leading-edge annotations as inferred by vector field integration under baseline and INHBA-reduced conditions.

**Supplementary Figure 5:**
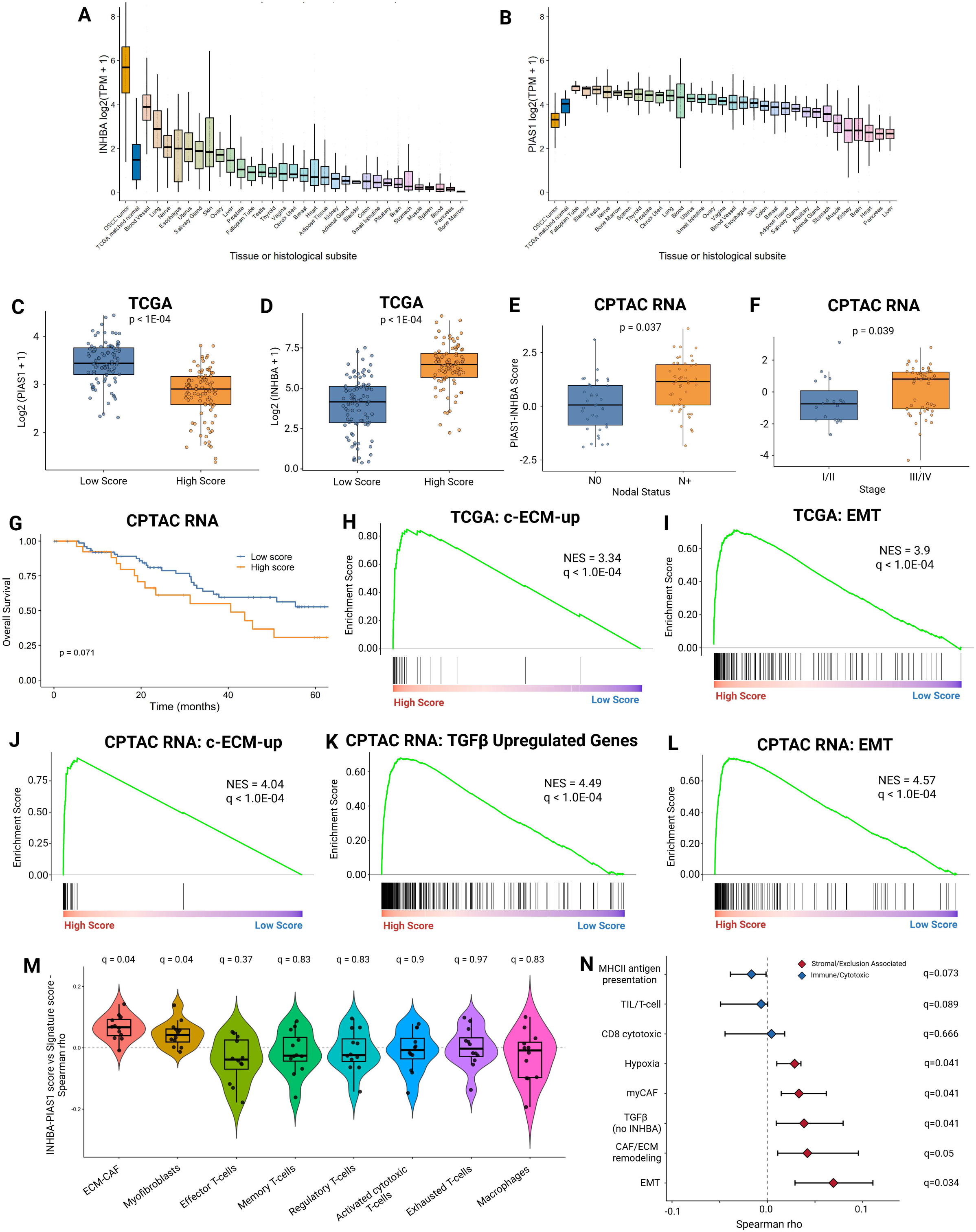
INHBA-PIAS1 score analyses across normal tissues, TCGA, and CPTAC cohorts. (A) Box plots showing INHBA expression across OSCC and normal body subsites. (B) Box plots showing PIAS1 expression across OSCC and normal body subsites. (C) Box plot showing PIAS1 expression in low and high INHBA-PIAS1 score groups in the TCGA OSCC cohort. (D) Box plot showing INHBA expression in low and high INHBA-PIAS1 score groups in the TCGA OSCC cohort. (E) Box plot comparing the INHBA-PIAS1 RNA score in node-negative (N0) and node-positive (N+) tumors in the CPTAC cohort. (F) Box plot comparing the INHBA-PIAS1 RNA score in early-stage (I/II) and advanced-stage (III/IV) tumors in the CPTAC cohort. (G) Kaplan-Meier curve for OS based on high and low INHBA-PIAS1 RNA score groups in the CPTAC cohort. (H-I) Gene set enrichment plots showing enrichment of C-ECM-up and EMT programs in high versus low INHBA-PIAS1 score tumors using TCGA RNA data. (J-L) Gene set enrichment plots showing enrichment of C-ECM-up, TGFβ-upregulated genes, and EMT programs in high versus low INHBA-PIAS1 score tumors using CPTAC RNA data. (M) Violin plots showing Spearman correlations between the INHBA-PIAS1 score and indicated cell-type signature scores in spatial transcriptomics data. (N) Forest plot showing Spearman correlations between the INHBA-PIAS1 score and immune, stromal, hypoxia, TGFβ, CAF/ECM remodeling, and EMT-related signatures in spatial transcriptomics data.

**Supplementary Table 1:** Clinicopathological characteristics of Calgary cohort (n=60), including demographic, tumor staging, and lifestyle factors.

| Variable | n (%) unless otherwise indicated |
| --- | --- |
| **Number** | 60 |
| **Sex** |  |
| *Female* | 15 (25) |
| *Male* | 45 (75) |
| **Median age at diagnosis, years (IQR)** | 62.5 (54.75 - 69.25) |
| **Pathologic T stage** |  |
| *T1/T2* | 16 (26.7) |
| *T3/T4* | 44 (73.3) |
| **Pathologic N stage** |  |
| *Node-negative* | 26 (43.3) |
| *Node-positive* | 34 (56.7) |
| **Overall Stage** |  |
| *Stage I/II* | 11 (18.3) |
| *Stage III/IV* | 49 (81.7) |

**Supplementary Table 2:** Clinicopathological characteristics of ORI OSCC TMA cohort (n=175), including demographic and tumor staging.

| Variable | n (%) unless otherwise indicated |
| --- | --- |
| **Number** | 175 |
| **Sex** |  |
| *Female* | 65 (37.1) |
| *Male* | 110 (62.9) |
| **Median age at diagnosis, years (IQR)** | 62.5 (53.9 - 74.8) |
| **Pathologic T stage** |  |
| *T1/T2* | 105 (60.0) |
| *T3/T4* | 62 (35.4) |
| *Missing* | 8 (4.6) |
| **Pathologic N stage** |  |
| *Node-negative* | 63 (36.0) |
| *Node-positive* | 79 (45.1) |
| *Missing* | 33 (18.9) |
| **Overall Stage** |  |
| *Stage I/II* | 61 (34.9) |
| *Stage III/IV* | 107 (61.1) |
| *Missing* | 7 (4.0) |

**Supplementary Table 3:**
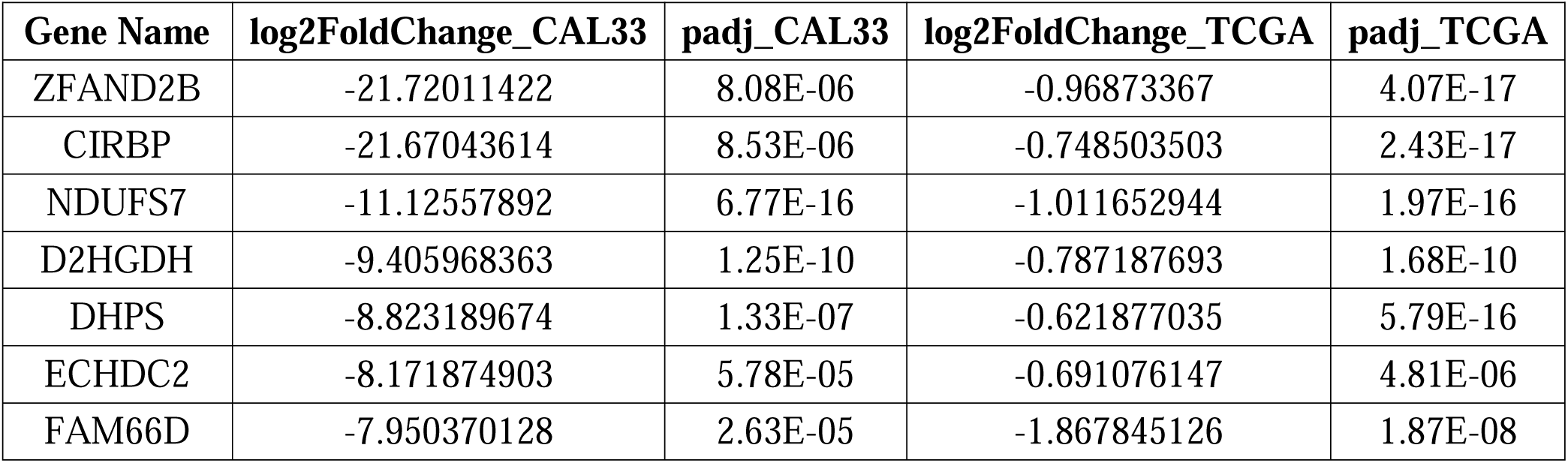
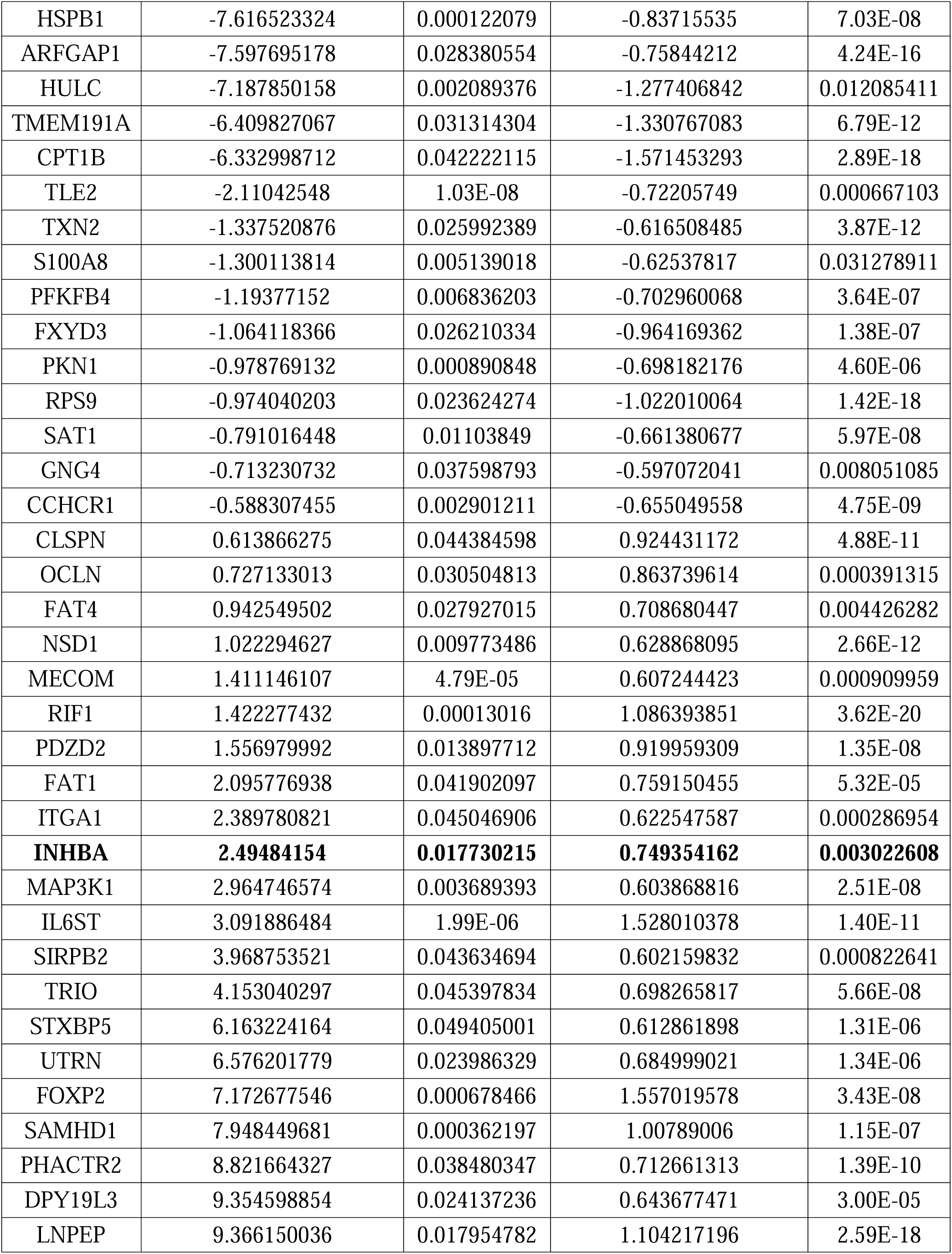
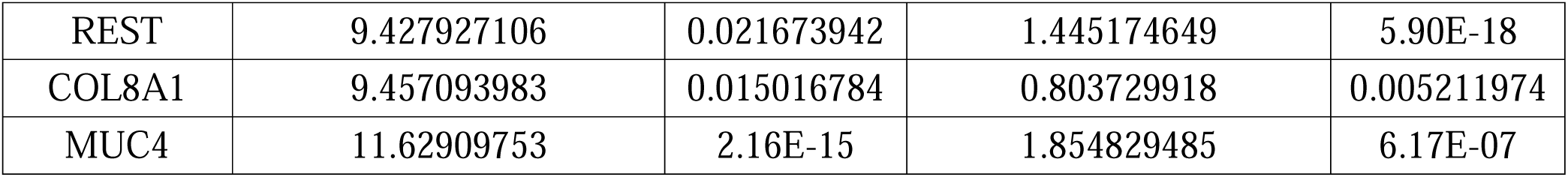
Overlapping genes and their fold change (log2 scale) and adjusted p-value (padj) in CAL33 PIAS1i and TCGA. INHBA is highlighted.

