## Supplementary figures and images for "PIAS1 loss and INHBA gain define a TGFβ-driven EMT state in OSCC progression"

### Full Blots

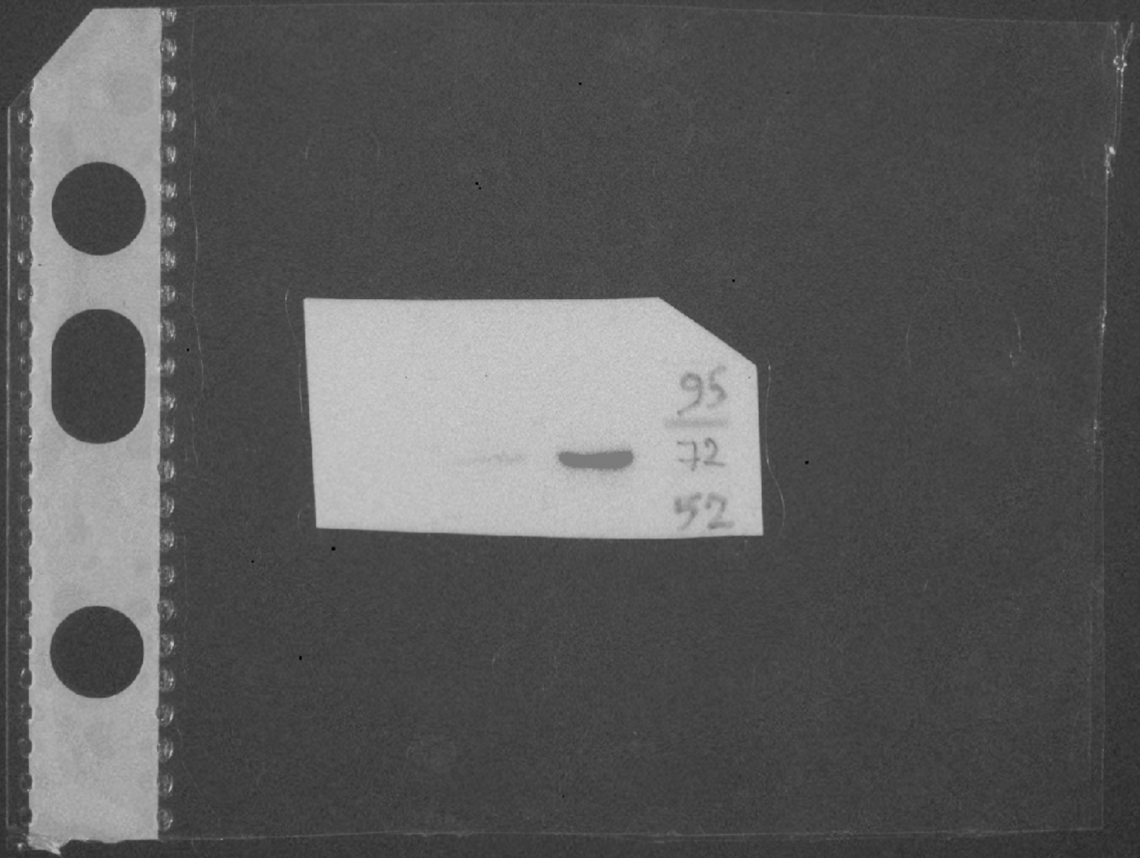

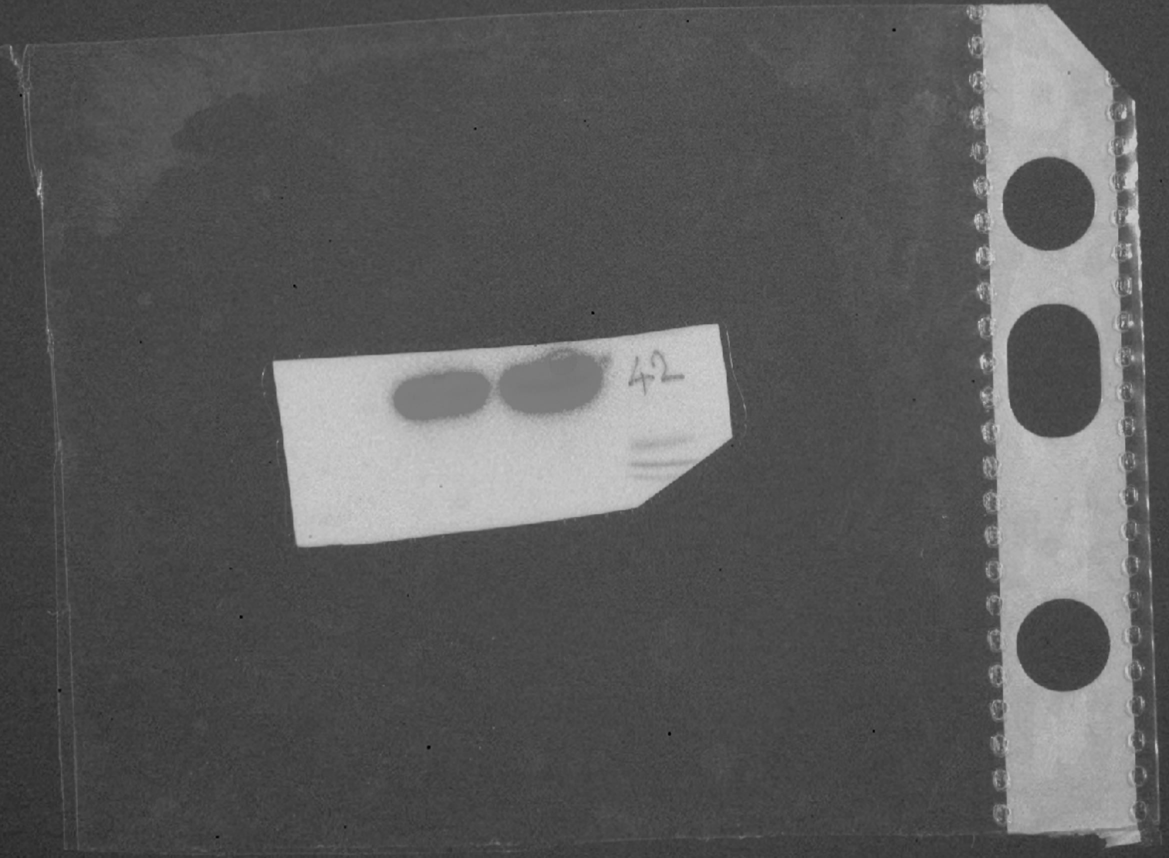

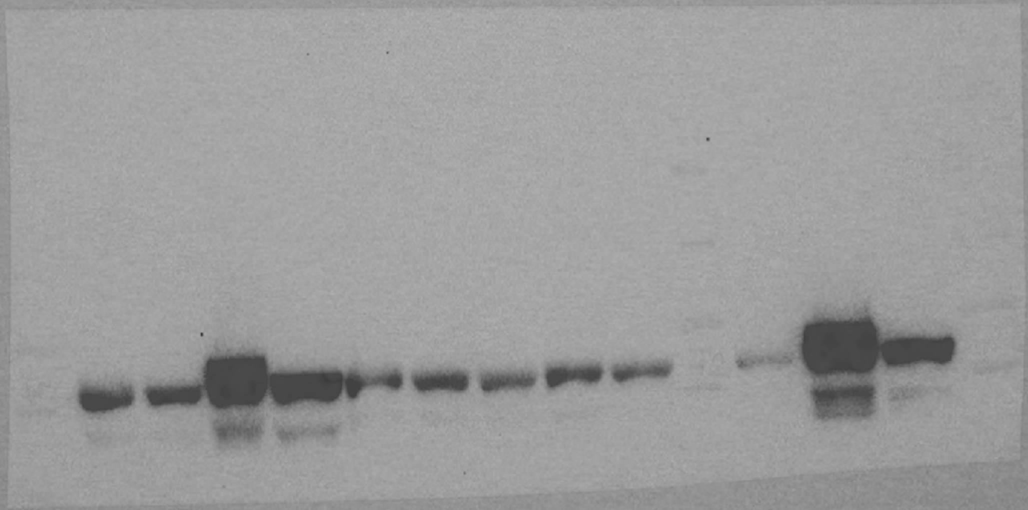

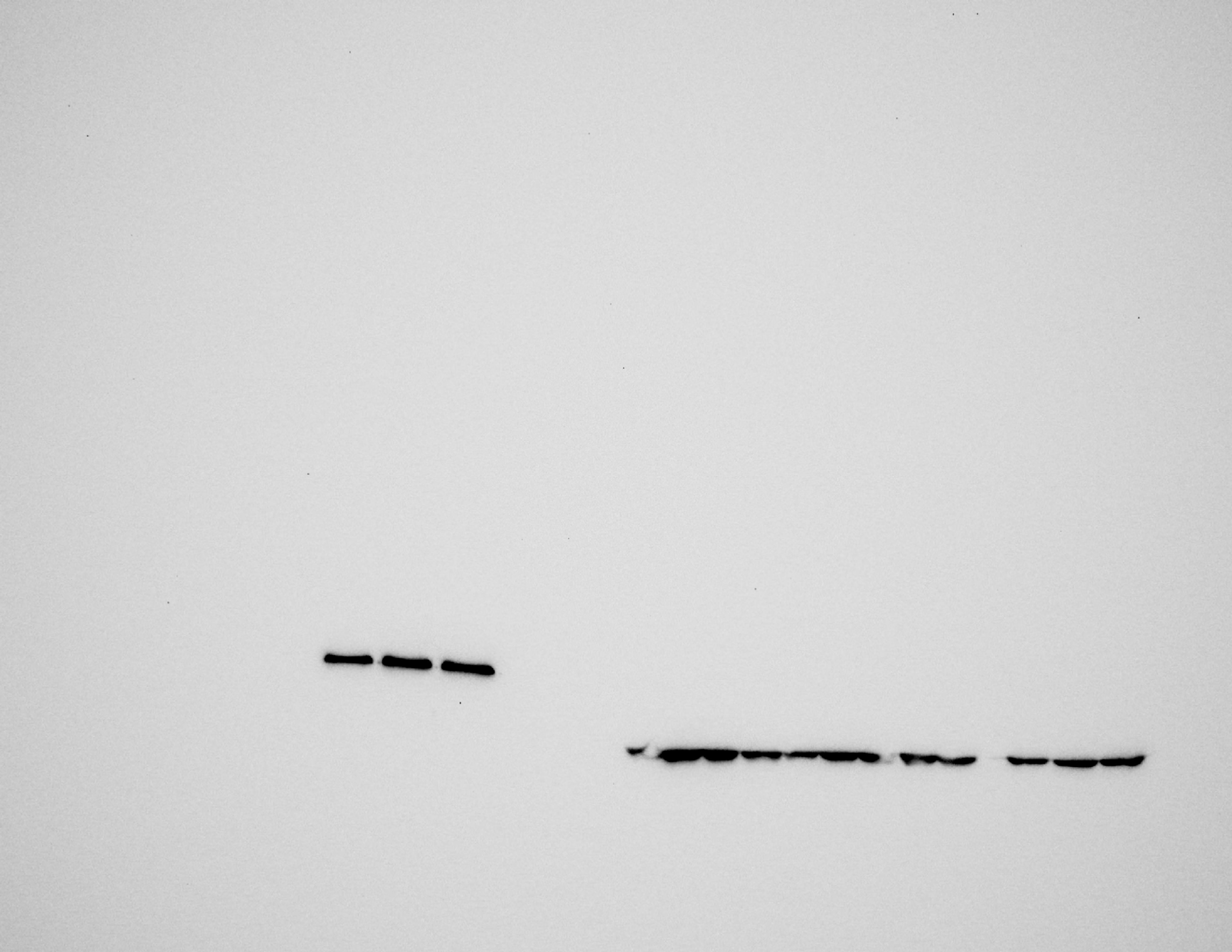
